# Mechanism of dopamine traveling waves in the striatum: theory and experiment

**DOI:** 10.1101/2022.04.19.488647

**Authors:** Lior Matityahu, Naomi Gilin, Yara Atamna, Lior Tiroshi, Jeffery R. Wickens, Joshua A. Goldberg

**Author notes:** Department of Neurobiology, Northwestern University, Evanston, IL, 60208, USA.

## Abstract

Striatal dopamine (DA) encodes reward, with recent work showing that DA release occurs in spatiotemporal waves. However, the mechanism of DA waves is unknown. Here we report that the striatal cholinergic neuropil also exhibits wave-like activity, and that the spatial scale of striatal DA release is extended by nicotinic receptors. Based on these findings we hypothesized that the local reciprocal interaction between cholinergic interneurons (CIN) and DA axons suffices to drive endogenous traveling waves. We show that the morphological and physiological properties of the CIN-DA interaction can be modeled as a reaction-diffusion system that gives rise to traveling waves. Analytically-tractable versions of the model show that the structure and the nature of propagation of CIN and DA traveling waves depend on their coupling, and that traveling waves can give rise to empirically observed correlations between these signals. Our model provides a biophysical mechanism for wave formation and predicts that the observed DA and CIN waves are strongly coupled phenomena.

## Introduction

Striatal dopamine (DA) is essential for motivated behavior and reinforcement learning. Consistent with a role in these processes, the activity of DA neurons has been associated with reward prediction error^1, 2^ and other motivationally significant events^3^. Clinically, degeneration of DA neurons causes Parkinson’s disease and striatal DA depletion causes motor symptoms^4^. To understand how DA contributes to these functions the nature of DA neurotransmission has been extensively studied. The extensive arborization of DA axons^5^, the high density of release sites^6^, and the location of DA receptors and transporters some distance from release sites has led to the concept of DA as a volume-transmitted, global and spatially-uniform signal^7^. In contrast to the concept of a spatially-uniform signal, a recent imaging study has shown spatiotemporal traveling waves of DA^8^. These traveling waves of DA concentration traverse the mediolateral (ML) aspect of the striatum, and are evident both in DA dynamics imaged using the genetically encoded DA sensor, dLight^9^ and in DA axon activity, imaged using genetically encoded Ca^2+^ indicators (GECIs). The functional significance of these waves is suggested by evidence that medial to lateral waves were associated with instrumental learning, while lateral to medial waves were associated with the reward delivery during classical conditioning^8^. How these waves arise and travel is currently unknown. Here we investigate the dynamical mechanism that gives rise to the formation and propagation of striatal traveling DA waves.

Studies indicating local modulation of DA release in the striatum provide an intriguing clue to the underlying mechanisms of striatal traveling DA waves. Several studies have shown gradual increases of DA concentration – DA ramps – during cued reward^10, 11, 12, 13, 14^. These gradual ramps differ from the transient, phasic increases in firing activity of DA cell bodies in the midbrain recorded during similar cued reward tasks^15^ or locomotor acceleration^16^. Although the encoding function of DA ramps is hotly debated^17^, there is mounting evidence that DA release is locally modulated, independent of DA cell firing at the soma^10, 11, 12, 13, 18, 19, 20^.

Here we investigate the hypothesis that the traveling waves of dopamine are generated locally in the striatum. This hypothesis of a striatal origin of the DA waves is based on two observations. First, several studies have shown that striatal cholinergic interneurons (CINs) modulate the release of DA by actions at nicotinic receptors on DA terminals^21, 22, 23, 24^. Striatal DA axons express α4β2 nicotinic acetylcholine receptors (nAChRs), whose activation can “hijack” the axons and lead to local DA release^20^. Activation of striatal CINs causes striatal DA release *in vitro*^21, 22, 23^. Thus, CINs exert local control over DA release and DA-mediated behavior^25, 26^.

Second, we previously found preliminary indications of wavelike dynamics – observed with GECIs – in the neuropil of striatal cholinergic interneurons (CINs) in freely moving mice^27, 28^. The putative coexistence of acetylcholine (ACh) and DA waves in the striatum suggests a possible coupling of these two phenomena. Various studies have shown that CIN signaling is time-locked to DA signaling in the striatum, with some studies finding an anti-phase relationship between the two signals while others found out-of-phase coupling^19, 29, 30^. Moreover, the correlation structure may depend on whether, for example, a cue or reward is being presented^8, 29^. The putative presence of coincident CIN and DA traveling waves will dictate a particular correlation structure between these two signals.

In the current study, we first present novel evidence of wave activity in CIN neuropil in freely moving mice. We then report that nAChRs extend the distance over which DA release can be detected after electrical stimulation by several hundred micrometers. Based on these findings, we propose a dynamical scheme by which the local *reciprocal* interaction between CINs and DA fibers gives rise to locally generated traveling waves of both DA and CIN activity, and cause temporal correlations that are similar to those observed empirically. We will also discuss parameter regimes where this interaction between DA and CINs can give rise to the formation of spatial Turing patterns^31^ manifest as “hills of activity” of DA and ACh, that may dynamically parcel the striatum into distinct functional regions of high vs. low concentrations of these two neuromodulators.

## Results

### Wave-like activity in the striatal cholinergic neuropil

We have previously shown that activity within the cholinergic neuropil of the striatum, visualized microendoscopically with the GECI GCaMP6s in freely moving mice, is highly synchronized across the striatum and acts as a measure of collective CIN activity ^27, 28^. At times we observed directional spreading of the GCaMP6s signal through the neuropil. In light of the recent finding of striatal DA waves, we re-analyzed those data, using optic-flow techniques that were implemented in the analysis of DA waves^8^. Our re-analysis corroborated the existence of directional flow of cholinergic activity in the striatum of freely moving mice (Fig. 1, Suppl. Movie 1). The fact that both signals – CIN activity and DA release – exhibit wave-like activity raises the possibility that these two signals are coupled, and may be generated by a joint mechanism, which is the central hypothesis of the current study.

**Figure 1.**
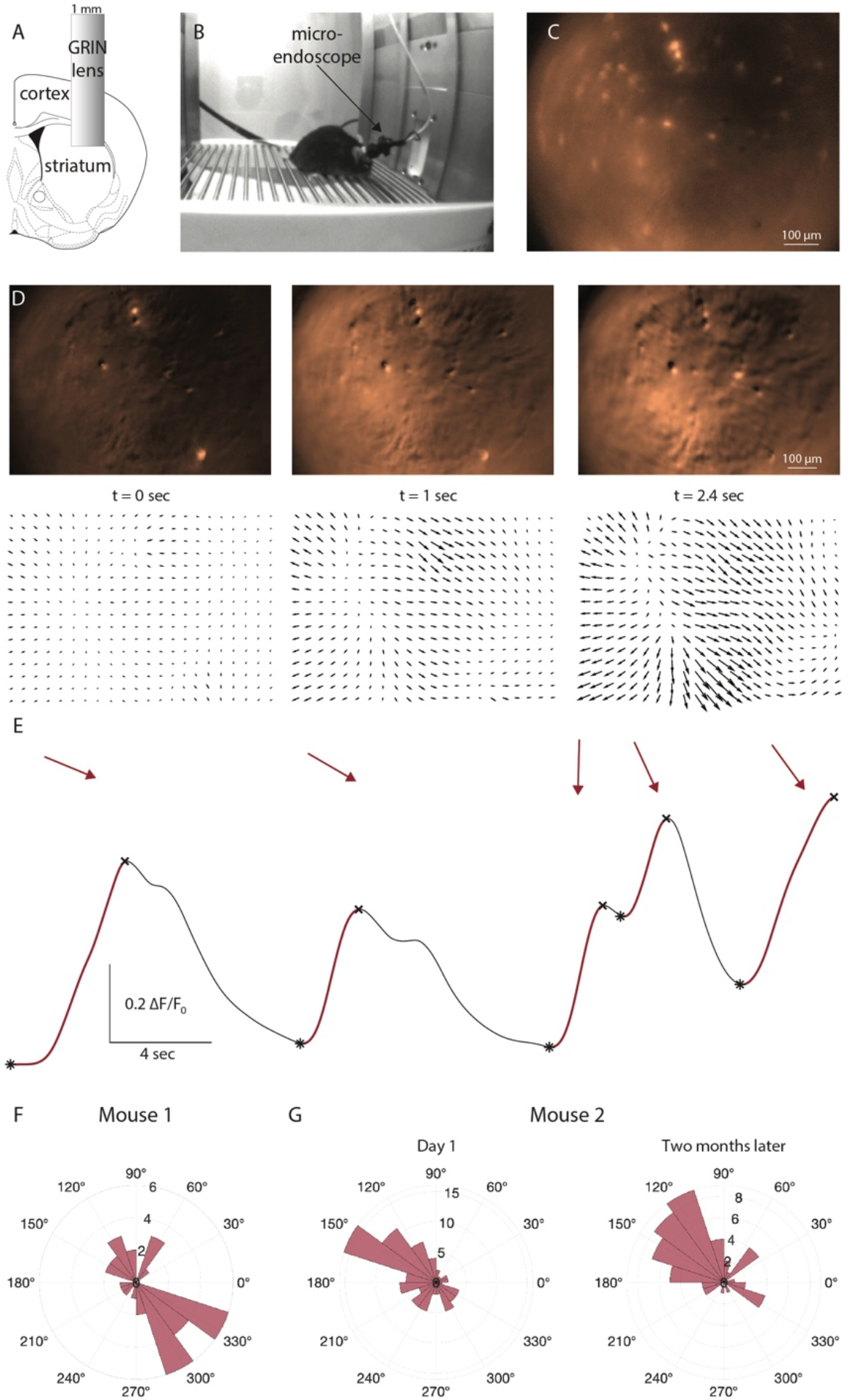
Wave-like activity in striatal cholinergic neuropil in freely moving mice. **A.** A 1 mm diameter GRIN lens is implanted into dorsolateral striatum following aspiration of cortical tissue and viral inoculation. **B.** Implanted mouse with a microendoscope mounted on its head moves freely in a behavior chamber. **C.** Image via GRIN lens in freely moving mouse reveals the raw GCaMP6s signal from several dozen identifiable somata and from the surrounding neuropil. **D.** Isolation of the microendoscopic background signal using the CNMF-E algorithm and calculation of Δ*F/F_0_* for this signal (see Methods) revealed events of elevated fluorescence (top row) that expanded to the bottom right of the field-of-view, as revealed by an optic flow algorithm^8^. **E.** To analyze the reproducibility of these events we calculated the spatial average of the background signal as a function of time (black trace) and isolated the large upswings in this signal (maroon trace). For each upswing we averaged over the duration of the upswing the average spatial vector (maroon arrows above the upswing, drawn in the same orientation of the figures in D) from all the pixels, whose associated flow vector was deemed significant using a bootstrapping method (see Methods). **F.** The angular histogram of all the average vectors measured throughout the microendoscopic imaging session exhibited a large degree of reproducibility in the directionality of the background neuropil signal. **G.** Same as a F, but for another mouse, with histograms for two recording session conducted two months apart.

### The spatial extent of striatal DA release is under the control of nAChRs

A necessary condition for the spread of CIN and DA activity to share a common mechanism, is that CIN and DA activity are coupled, and that the coupling contributes to the spreading *per se* of the activity. While it is known that CINs can activate nAChRs on DA axons to drive DA release^21, 22, 32^, we wanted to determine whether this activation also extends the range of DA release. We therefore expressed a genetically-encoded DA sensor, GRAB-DA2m, in striatal DA axons, and measured the spatial extent of DA release in response to electrical stimulation of an acute striatal slice (Fig. 2A). The electrical stimulation triggered DA release several hundred micrometers from the bi-polar electrode (Fig. 2B, “control”). Estimation of the spatial profile of release, demonstrated that it fell off with a spatial scale of approximately 500 μm (Fig. 2C, “control”). Application of 10 μM of the nAChR antagonists, mecamylamine (Fig. 2B, “mecamylamine”), halved the spatial scale of DA release (Fig. 2C, “mecamylamine”), suggesting that the more distal release of DA depended on the recruitment of CINs in the proximity of the electrode by the electrical stimulation. It is these CINs that presumably caused the distant DA release by activating nAChRs on DA axons. Measurement of the spatial scale of recruitment of CINs supported this conclusion. The spatial extent of the recruitment of CINs – estimated by measuring the fall off of the GCaMP6f signal in CINs in response to the same electrical stimulation – was estimated at approximately 200 μm (Fig. 2C, “ChAT-GCaMP6”). These findings suggest that activation of CINs in one region of the striatum lead to – or at least enhanced – the distant release of DA. We therefore conclude that local coupling between CINs and DA can cause the spread of the DA release.

**Figure 2.**
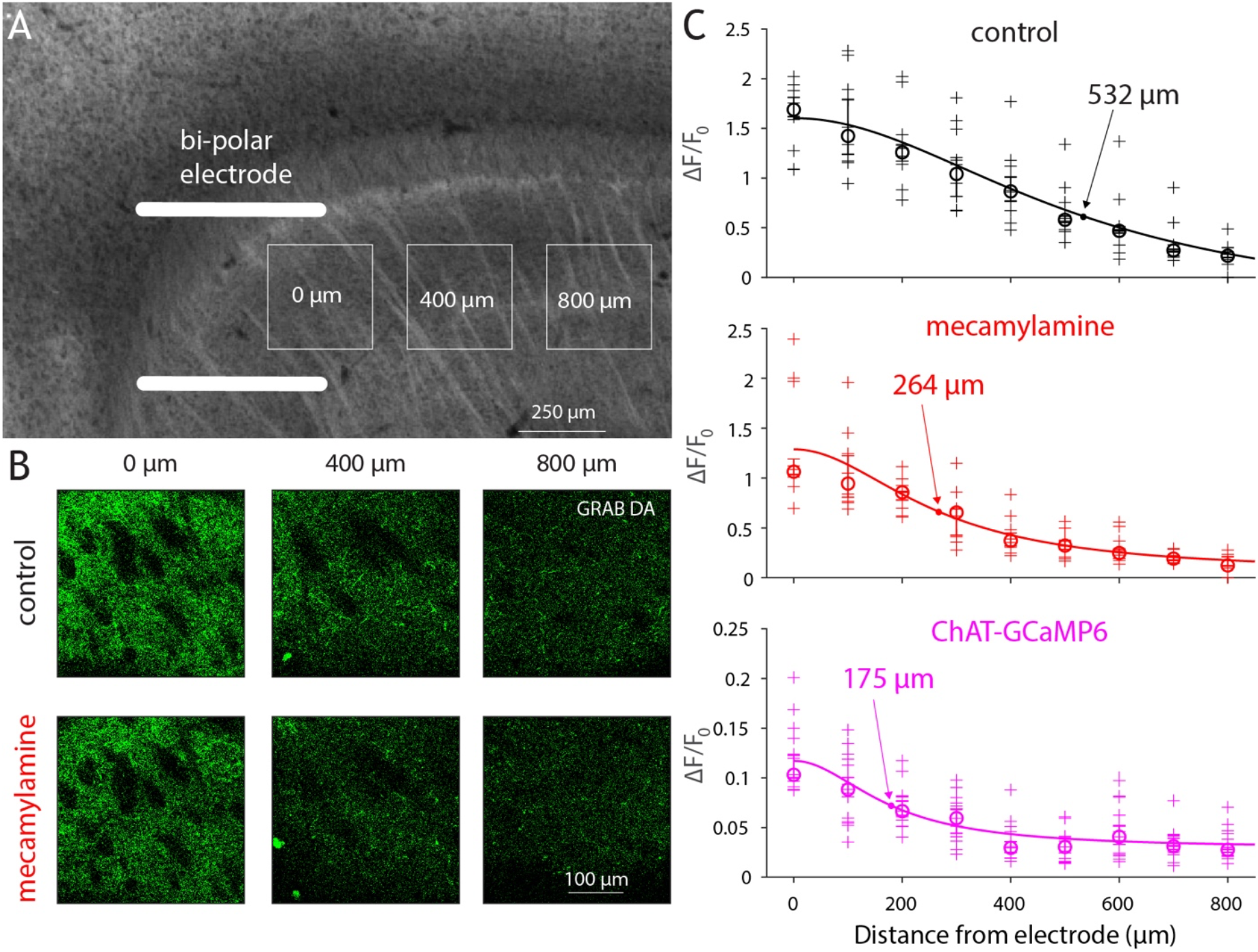
nAChRs increase the spatial extent of striatal dopamine release. **A.** Experimental design. Acute sagittal slice of striatum from mouse whose SNc was inoculated with AAVs harboring GRAB-DA2m is stimulated with a bi-polar electrode (drawn to scale). Maximal GRAB-DA2m fluorescence in response to electrical stimulation is measured within 300 μm × 300 μm ROIs, centered at 100 μm intervals from the electrode (0, 400 and 800 μm shown). **B.** Raw GRAB-DA2m fluorescence at various distances from the bipolar electrodes in control (top) and after bathing slice in 10 μM mecamylamine, an nAChR antagonist. **C.** Decay of GRAB-DA2m fluorescence (Δ*F/F_0_*) as a function of distance in control (top, *n* = 11 slices from *N* = 2 mice) and in mecamylamine (middle, ANCOVA, F(1,176) = 5.57, P = 0.0194), and decay of GCaMP6 fluorescence in cholinergic neuropil from ChAT-GCaMP6f mice as a function of distance (bottom, *n* = 15 slices from *N* = 3 mice). Crosses – individual measurements; circles – median; Solid curve – fit of Lorentzian to data points, from which the indicated spatial constants were extracted (See Methods).

### An extended model of local coupling between CINs and DA

The finding that both DA release^8^ and CIN neuropil exhibit wave-like activity, in conjunction with our demonstration that CINs can extend the release of striatal dopamine, led us to hypothesize that it is the local coupling between CIN and DA axons that produces the wave-like activity. Moreover, we hypothesized that this coupling occurs throughout the densely intertwined arborization of CIN and DA processes^20^, causing the two neuromodulatory systems to behave like a nonlinear coupled reaction diffusion system that can give rise to traveling waves. We therefore constructed a reaction-diffusion model that replicates the main features of the known coupling between CINs and DA, as described presently.

The dynamical scheme of our model is that of an activator inhibitor reaction diffusion (AIRD) system, which is known in chemistry and biology to give rise to traveling wave phenomena as well as to spatial Turing patterns^31, 33^. As the name suggests, AIRD systems involve both activators and inhibitors. Because CINs activate nAChRs on DA axons – even spontaneously under certain *in vitro* conditions^20, 34, 35, 36^ – we assume that CINs are the activators in the system. Conversely, the DA fibers are the inhibitors because DA release inhibits CIN activity via DA D2 receptors (D2Rs)^20, 37^.

Modeling CIN-DA axon interactions as a reaction diffusion system is justified by the dense, tortuous, space-filling nature of both DA and CIN axons, which have release sites every few microns^20, 38^. This structure combined with the fact that DA and ACh may diffuse some distance from release sites, lends support to modeling them as continuous media (or syncytia) that fill space and interact at every point. Finally, the activators in AIRD systems need to be “autocatalytic” (i.e., self-exciting thorough positive feedback). At first sight, this seems highly improbable, because CINs are known to be mutually coupled by di-synaptic inhibition^39, 40^. Nevertheless, we will propose one mechanism by which CINs may be self-exciting, due to the non-monotonic dependence of nAChRs on ACh concentration, and discuss other possibilities, as well.

### Derivation of the coupled CIN-DA model

We model CINs using a threshold linear rate model, with *C* denoting their rate

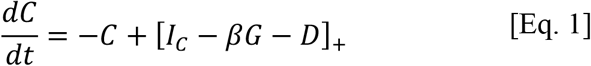

The autonomous firing of CINs^41^ is realized by a constant input current, *I_C_* > 0. *G* represents the firing rate of the intrinsic source of striatal inhibition, which arises *in vitro* exclusively from GABAergic interneurons (GINs)^42, 43, 44^ *β* > 0 represents the gain of the GIN-CIN connection, and [%]_+_ ≝ *max* (*x*, 0). *D* represents the DA that inhibits CINs via activation of DA D2 receptors (D2Rs). The integration time constant of *C* is assumed to be unity. The equation for *G* is given by

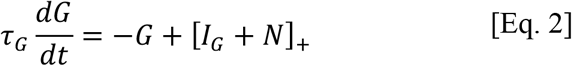

where *I_G_* > 0, gives rise to the tonic activity of the GINs^44, 45, 46^. *N* is the activation of the α4β2 nAChRs on the GINs. Note that here the threshold-linear gain function is superfluous and can be omitted. The integration time constant of the GINs is denoted by *τ_G_*. The equation for *N* is given by

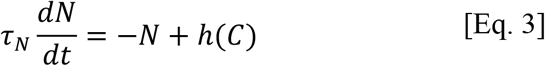

where the integration time constant of the nAChRs is denoted by *τ_N_* (Fig. 3A). The function *h*(·) represents the dependence of the α4β2 nAChRs activation on the activation of CINs. α4β2 nAChRs activation has an inverted-U shaped dependence on to concentration of ACh, and behaves roughly like 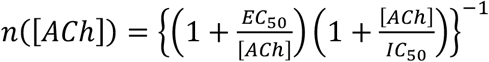 with EC_50_=175 μM and IC_50_=12 mM^47, 48^. Note that *n*(·) peaks in the 1 millimolar range. ACh is thought to reach concentrations of 1-10 mM at the post-synaptic end of the synaptic cleft of the neuromuscular junction^49^ and similar concentrations in the proximity of CIN release sites in the striatum^38^. Because of the wide range of concentrations, the inverted-U region of this function – namely where an increase in ACh concentration can lead to *less* activation of nAChRs – is physiologically relevant (Fig. 3B). Assuming that there is a monotonic (increasing) dependence of striatal ACh concentration on the activity of CINs, we will model *h*(·) as an inverted-U function, as well, namely *h*(*x*) ≥ *xe*^−*κx*^ (*κ*> 0).

**Figure 3.**
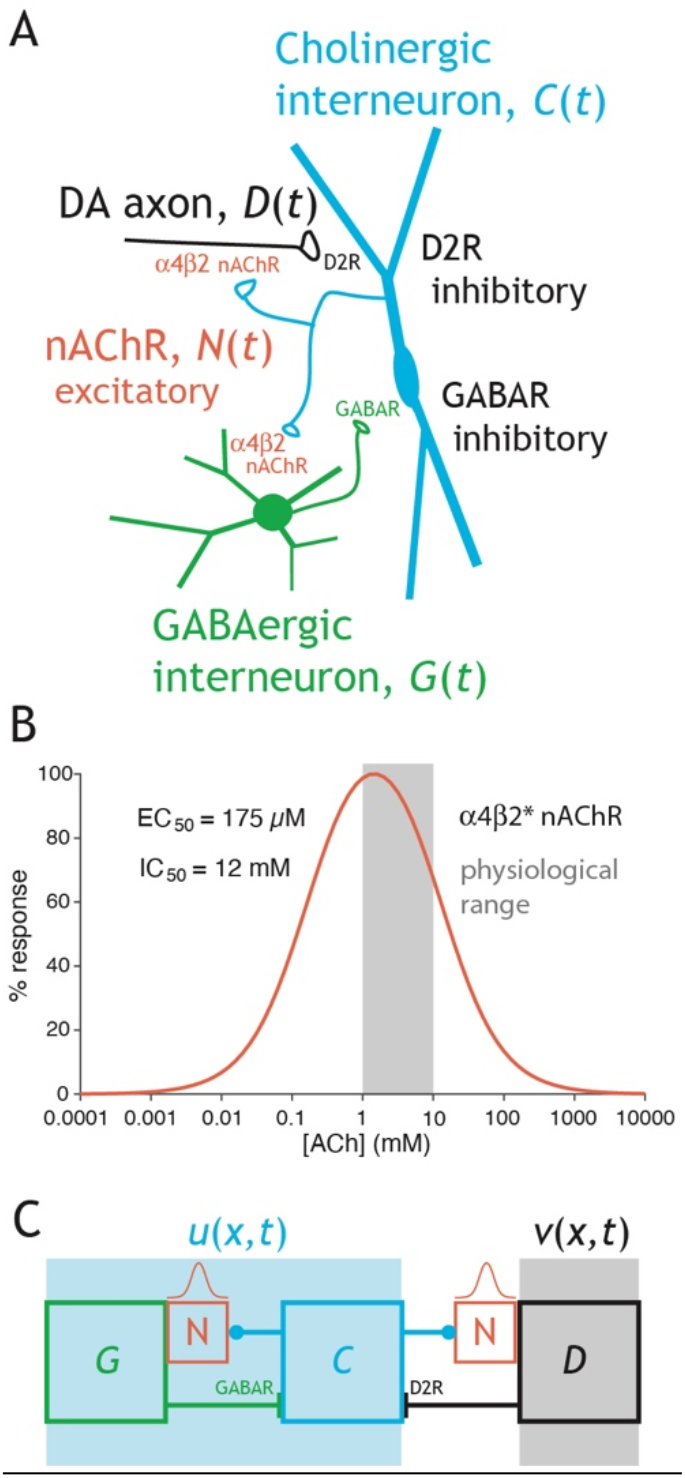
Activator-inhibitor reaction diffusion (AIRD) model of local striatal interaction between CINs and DA axons. **A.** Circuit diagram of the reciprocal interaction between CINs and DA axons, as well as between CINs and GINs, and the dynamical variables use to denote the various elements, including the nAChRs. **B.** Proposed inverted-U-shape dependence of nAChR activation on ACh concentration. The 1-10 μM range, indicated in gray, is hypothesized to be the physiological range, wherein an increase in concentration leads to a reduction in activation. **C.** Extended AIRD model where the reciprocal interaction between CINs and GINs is subsumed into the variable *u*(*x,t*), and the DA is represented by *ν*(*x,t*) (Eqs. 4-6). *ν* is governed by an inverted-U dependence on *u*, and *u* is governed by an inverted-N dependence on itself.

Because the kinetics of the nAChRs are faster than those of the neurons, we will assume Eq. 3 is at steady state. Similarly, we will assume the GINs’ response to activation of nAChRs is faster than the CINs response to activation by GINs or in other words that *τ_G_* ≪1. Therefore, we will use the steady state solution of Eq. 2, as well. Thus, taken together we can reduce the dynamics of the CINs to

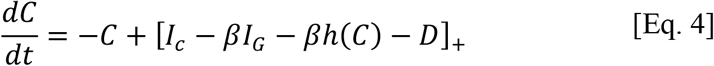

We will be interested in the parameter regime where the argument of the threshold-linear function is positive, hence Eq. 4 can be simplified to

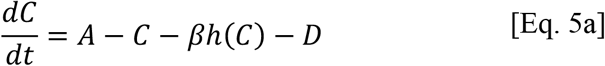

where *A* = *I_C_* – *βI_G_* (Fig. 3C, light blue background). The equation for the *D* is

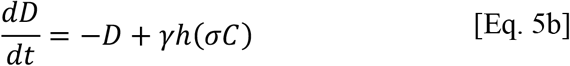

because DA is activated by the nAChRs which are activated by the CINs (Fig. 3C). *γ* > 0 represents the gain of the nicotinic innervation of the DA fibers, and *σ* > 0 is related to the “affinity” of the nAChRs on DA fibers to the activation of CIN (i.e., *σ* =*1* means that the nAChRs on GINs and on DA respond with identical sensitivity CIN activity).

As explained in the Introduction, we assume that the CINs can be represented by a spatially extended variable, *u*(*x,t*) (for simplicity we conduct the analysis in one dimension). DA is also represented as a spatially extended variable, *v*(*x,t*) (Fig. 3C). When diffusion is added, Eqs. 5 become the following coupled partial differential equations (PDEs) that describe the local coupling of CINs and DA fibers in the striatum

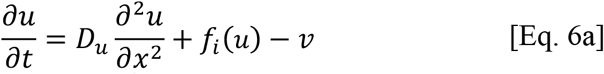

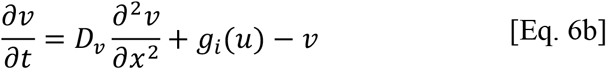

where *f*_1_(*u*) = *A* – *u* – *βh*(*u*) and *g*_1_(*u*) = *γh*(*σu*). We will refer to this model (for which the subscript *i*=*1*) as the “original model”. *D_u_* and *D_ν_* are the effective diffusion coefficient of *u* and *ν*, respectively. In this model, we are not considering physical diffusion of ACh and DA, but rather the “diffusion” of the activity. We will consider two regimes in our analysis. In the first we will assume that *D_ν_* = 0, which corresponds to a regime where activation of nAChRs on DA fibers, can induce local activation of these fibers (and presumably release of DA), but does not cause (electrical) activity to propagate throughout the DA axonal arbor.

In order to facilitate the analysis of Eqs. 6, we will also consider polynomial functions in place of *f_1_*(*u*) and *g_1_*(*u*) that will preserve the local geometry of the nullclines in the (*u,v*) phase plane. All simulations were run on XPPAUT^50^.

### Simultaneous advancing and receding traveling waves of CIN and DA activity

Traveling wave solutions in an extended medium arise when the dynamics of the medium are bistable such that one region of the medium is at one stable fixed-point solution of the diffusion-less system (i.e., *D_u_* = *D_v_* = 0 in Eqs. 6) while the other is at the other stable fixed-point. Then, when diffusion is re-instated, a traveling wave can form as a (moving) boundary between these two solutions. The velocity (direction and speed) of the wave is determined by the parameters of the equations that will determine which fixed point will eventually win-over the media (the wave solution and its direction will depend on initial conditions, as well). Bi-stability arises in Eqs. 6 due to the inverted-N shaped of *f*_1_(*u*) = *A* – *u* – *βh*(*u*) which results from the inverted-U shape of *h*(*u*) (Fig. 4). The two stable fixed points can be identified by the flow field around them. We will consider a regime where the nAChRs on DA fibers and GINs exhibit a similar sensitivity to ACh concentration (i.e., *σ* is close to unity). Note that in this region the two-stable fixed-points of the dynamics – in the absence of diffusion – are arranged geometrically such that the left stable fixed-point, *(u_1_, v_1_*), is a state of high DA and low CIN, whereas the right stable fixed-point, (*u_3_, v_3_*), is that of low DA and high CIN (Fig. 4A). In this case, when diffusion is introduced, the traveling wavefront of one variable advances while the other one recedes. The velocities of both wavefronts are equal, and depend on the parameters of *f_1_*(*u*) and *g_1_*(*u*) and on the diffusion coefficients. For example, increasing *β,* which represents increasing the gain of the nAChRs on the GINs, causes the CIN profile to transition from expanding to receding (Fig. 4B, Suppl. Fig. 2A).

**Figure 4.**
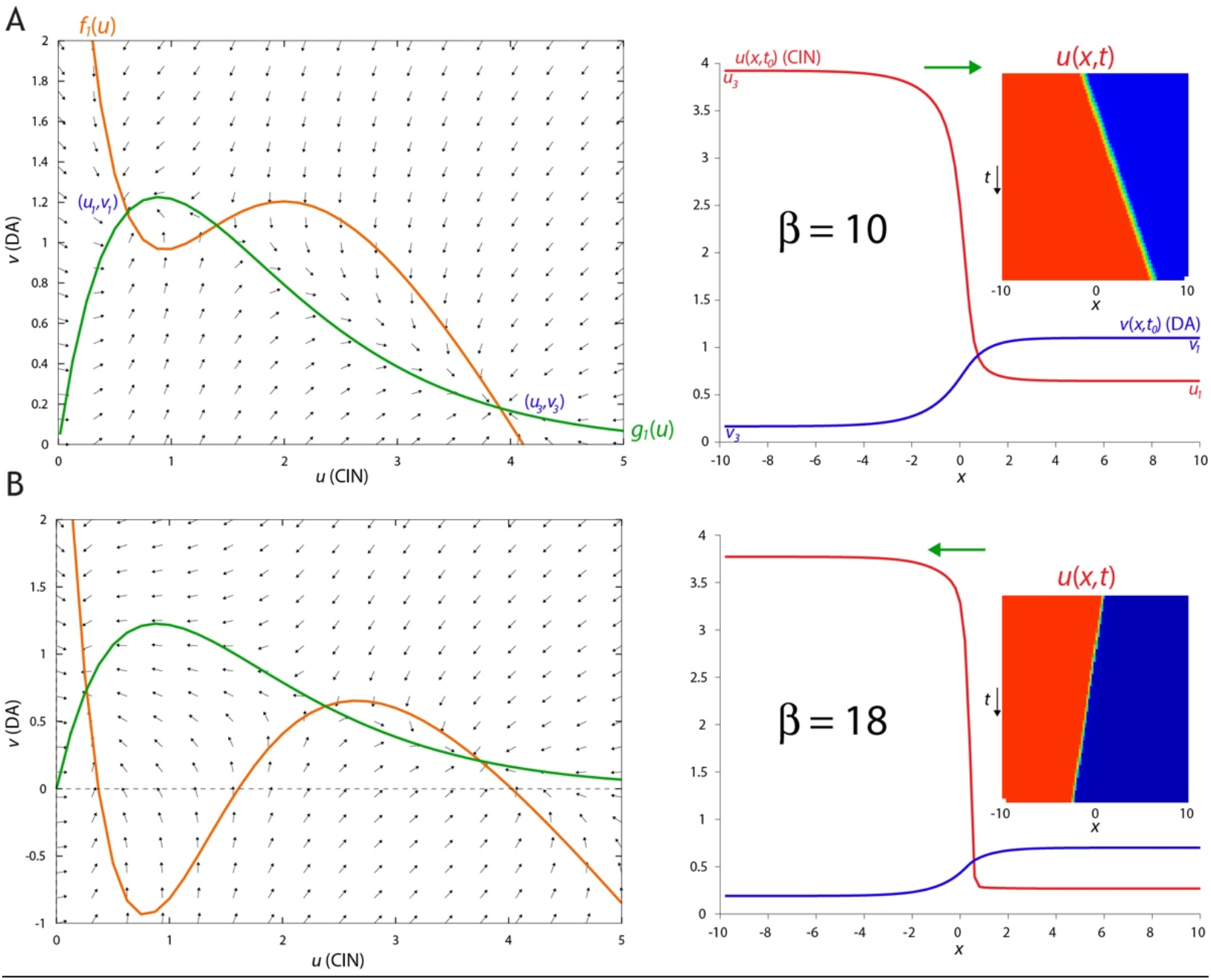
Simultaneous advancing and receding waves of DA and CIN activity in AIRD model. **A.** Left: Phase plane. The diffusion-less system’s nullclines intersect at 3 fixed-points, such that (*u_1_,v_1_*) and (*u_3_,v_3_*) (where *u_3_* > *u_1_* and *ν_3_* < *ν_1_*) are stable, as is evident from the flow fields. When *β* = 10 the area between the nullclines that is adjacent to (*u_1_*,*v_1_*) is smaller than the area that is adjacent to (*u_3_, v_3_*). Right: As a result the traveling waves of *u*(*x,t*) and *v*(*x,t*) move such that (*u_1_,v_1_*) wins over, meaning that the *u* (CIN) wave advances and the *v* (DA) wave recedes. Inset: space-time plot of *u*(*x,t*). **B.** Same as A, except that when *β* = 18, the area adjacent to (*u_1_,ν_1_*) is larger and the traveling waves are such that the *u* wave recedes and the *v* wave advances. Other parameters: *A=4.2, σ=0.75, κ=1.5, γ=4.7, D_u_=0.02, D_v_=1*.

To gain a fuller understanding of the behavior of the system, we will consider an analytically tractable version of Eqs. 6. In this case, *f_2_*(*u*) = *u*(1 – *u*)(*u* – *s*) + *a*^2^(1 – *u*) is an inverted-N shaped third-degree polynomial, and *g_2_*(*u*) = *bu*(1 –*u*) is an inverted-U shaped second-degree polynomial (Fig. 5A). In this model, we define *a_m_* = (*s* + *b*)/2 and require that 0 < *a* < *a_m_* < 0.5, which guarantees that, in the absence of diffusion, the system’s two stable fixed-points are

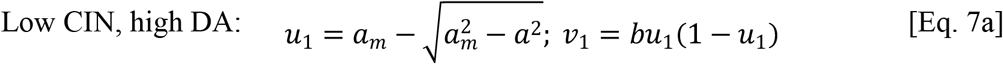

and

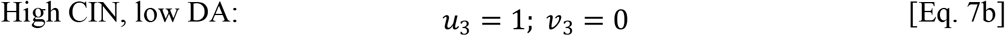

Thus, the purpose of the parameter *a* in *f_2_*(*u*) is to create the high DA solution (because *ν_1_* > *ν_3_* if and only if *a*>0). In this case, we search for a traveling wave solution for Eqs. 6, with velocity *c*, of the form

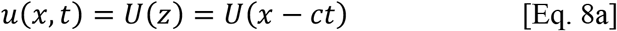

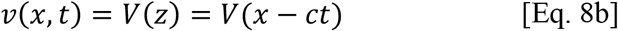

In this case, the PDE system (Eqs. 6) transforms into a pair of ordinary differential equations

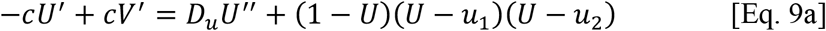

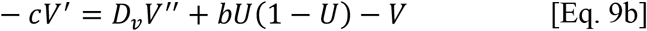

where 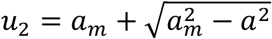, and the primes denote differentiation with respect to *z*.

**Figure 5.**
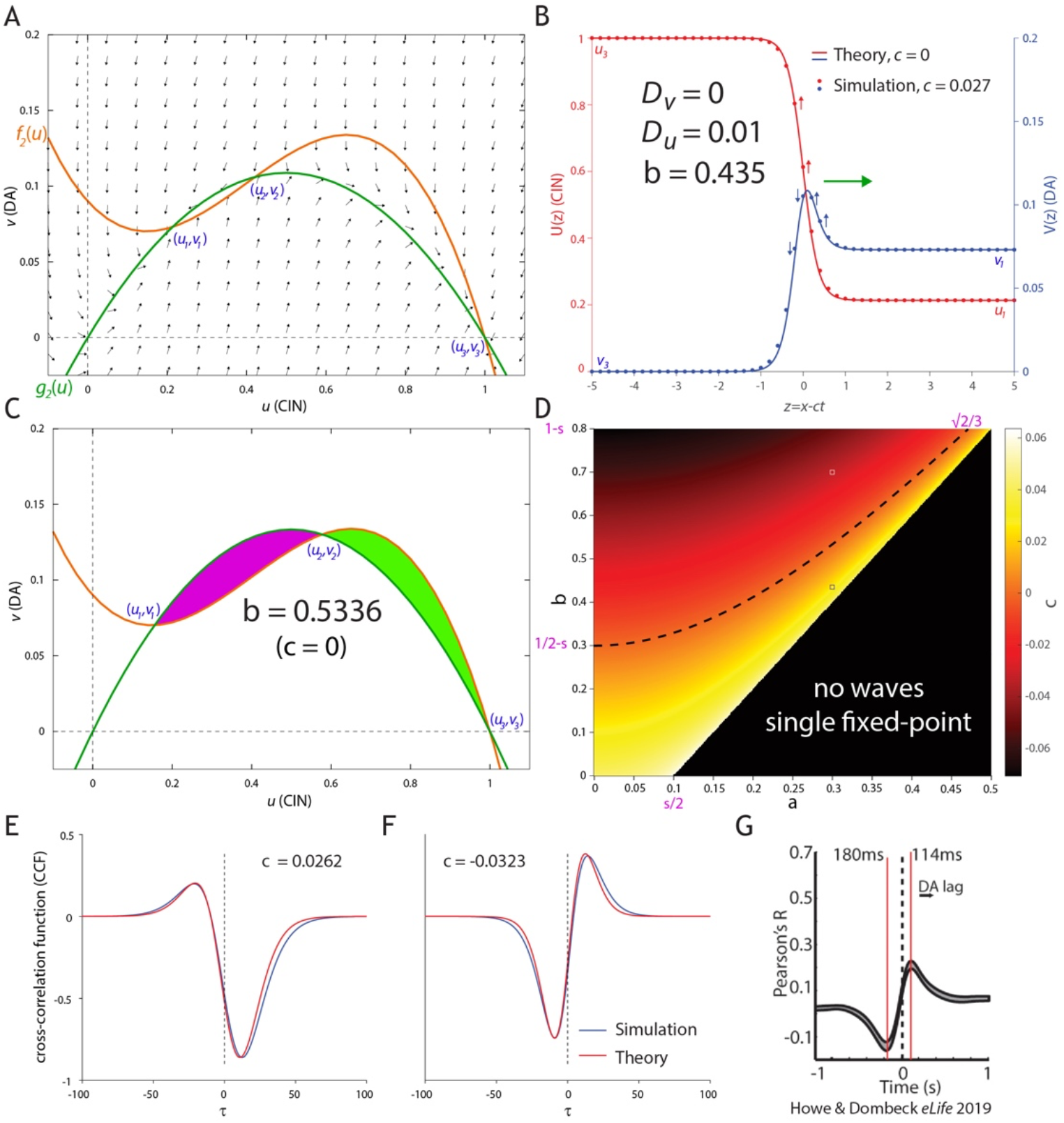
Tractable AIRD model produces various temporal correlation structures between DA and CIN depending on the structure and direction of the traveling waves. **A.** Left: Phase plane. The diffusion-less system’s nullclines intersect at 3 fixed-points, such that (*u_1_*, *v_1_*) and (*u_3_, v_3_*) (where *u_3_* > *u_1_* and *ν_3_* < *ν_1_*) are stable, as is evident from the flow fields. **B.** When diffusion of *u* is introduced (using the same parameters as in A), traveling waves form such that *u* advances and *ν* recedes, with velocity *c* that depends on these parameters according to Eq. 11. Analytical solution of waveform for *c* = *0* matches the simulated shape for small magnitudes of *c*. **C.** A standing wave of *c* = *0* forms for parameters where the (purple and green) areas adjacent to the two stable fixed-points are equal. **D.** Phase diagram. Dotted line indicates *c* = 0. Velocity according to Eq. 11 is color coded. **E.** Cross-correlation Function (CCF) between *u*(*x_0,t_*) and *v*(*x_0,t_*) as the traveling waves traverse *x_0_* for a positive value of *c* [blue – simulation; red – theory (Eq. 14)] for parameters corresponding to the white circle in D. **F.** Same as E, except that the value of *c* is negative and corresponds to the black circle in D. **G.** Reproduction of a recent empirical CCF – between CGaMP6 signals recorded with fiber photometry from DA and CIN striatal neuropil^19^ – that resembles the shape of the CCF in F. Other parameters: *a*=*0.3, s*=*0.2* (unless stated otherwise).

We can gain insight into the behavior of this system by considering the case of *D_ν_* = *0* (which corresponds to conditions where DA fibers can be locally activated by ACh, but this activation cannot propagate throughout the DA fibers). In this case, we can analytically solve the spatial profile of a standing wave (i.e., a traveling wave with velocity *c* = *0*), which is given by (Fig. 5B)

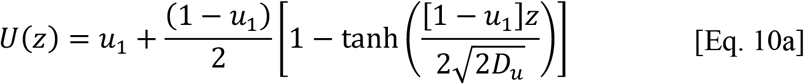

and

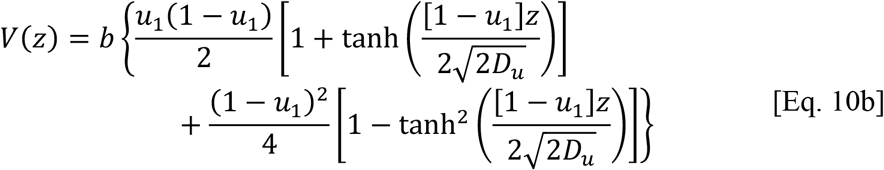

with the condition that *c = 0*, where

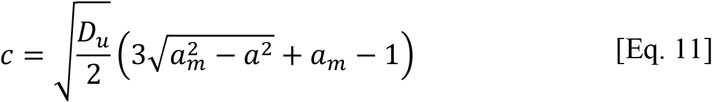

The overshoot in the profile of *V*(*z*), results from the fact that when *c* = 0, Eq. 9b can be rewritten simply as *V* = *bU*(1-*U*), so that the parametric trajectory (*U*(*z*), *V*(*z*)) runs along the green curve in Fig. 5A. Following the curve from (*u_3_,v_3_*) to *(u_1_,v_1_*), demonstrates that it overshoots the value of *v_1_* before approaching (*u_1_,v_1_*). Similar waveforms and traveling wave behavior can occur for *u*(*x,t*) and *v*(*x,t*) in the original model with *f_1_*(*u*) and *g_1_*(*u*) when *D_v_* = 0 (Suppl. Fig. 2B).

Note that the condition 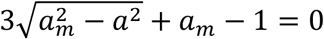 can also be derived by requiring that the integral 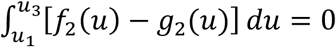, which is a general result for standing waves in bistable systems^51, 52, 53^. Geometrically this means that the areas confined by the two nullclines between each of the stable fixed points and the unstable fixed-point (*u*_2_) need to be equal to each other (purple and green areas in Fig. 5C). The condition *c* = *0* can be rewritten as the curve (Fig. 5D, dashed line)

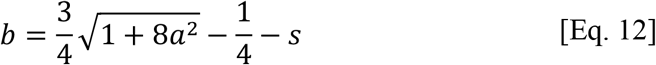

which splits the system’s phase diagram (Fig. 5D) into two regions: one (bottom right in Fig. 5D) where the *u* wave advances and *ν* wave recedes (i.e. the uniform solution, Eq. 7b, wins out); and the other (top left in Fig. 5D) where *u* recedes and *ν* advances (i.e. the uniform solution, Eq. 7a, wins out) (Suppl. Fig. 3). The values of the function *c*(*a,b;s,D_u_*) (Eq. 11) are color coded on the *b*-*a* phase diagram, even though they do not, in general, dictate the correct velocity for each point in the phase diagram. However, for small absolute values of *c* (and particularly if *b* ≪ 1) the term *cV*’ in Eqs. 9 will be negligible, so that the solution described in Eqs. 10 will be still approximately valid (Fig. 5B, “Theory” vs. “Simulation”).

### Temporal relationship between DA and CIN activity

A standard experimental method to characterize the temporal relationship between two signals is to calculate the temporal cross-correlation function (CCF) between them. If we consider any point *x_0_*, we can intuit how the two signals will change there in time. As the waveforms move to the right, i.e. *c* > 0 (Fig. 5B), the points along the wavefronts will change as indicated in the arrows (and this can be viewed directly in Suppl. Fig. 3). Formally, this means that that the change in activity at that point is given by the temporal derivative of the signals, i.e., –*cU*’(*x_0_*–*ct*) and –*cV*’(*x_0_*–*ct*). Thus, the CCF can be calculated as

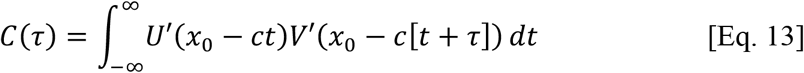

which, from translational symmetry of the traveling wave, is independent of *x_0_*.

If we consider *x_0_* = *0* in Fig. 5B, as the CIN signal there increases (upward red arrows), it will be positively correlated with the DA signals that are to its right (upward blue arrows). This means that the DA signal leads the CIN signal and the maximal correlation will be attained at a negative time delay, where DA precedes CINs (e.g., Fig 5E, blue curve). Note that in this case, there is a region at the leading edge of wavefront where the CIN and DA activities are elevated together. This region represents the ability of CINs to drive DA release by activating nAChRs on DA axons (Fig. 2)^20, 21, 22, 32^. Conversely, if the wavefronts move to the left, i.e., *c* < *0*, all the arrows will point in the opposite direction. In this case, the downward change in the CIN signal will be positively correlated with the lagging DA signals, which means that the maximal correlation will be attained at a positive delay, where DA lags behind CIN (Fig. 5F, blue curve).

The CCF can be calculated analytically for the wave solutions in Eq. 10. Because *V*(*z*) (Eq. 10b) is composed of two terms, the CCF is also composed of two terms, one symmetric, *C_S_*(*τ*), and another anti-symmetric, *C_A_*(*τ*):

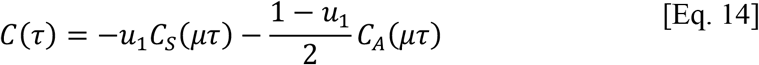

where

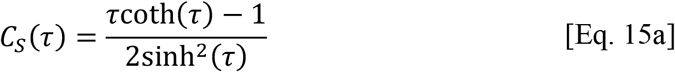

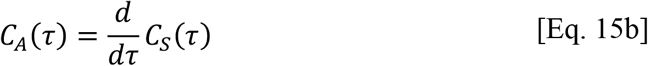

and

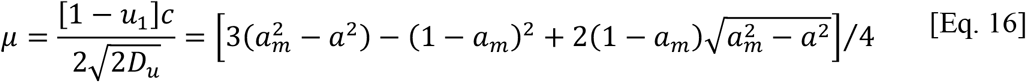

The theoretical calculation (Figs. 5E,F, red curves) closely resembles the numerical solution (blue curves). Intriguingly, the case where the CIN waveform recedes generates a CCF that resembles the empirical CCF measured recently between Ca^2+^ signals from CINs and striatal DA fibers that expressed GECIs, and were recorded with fiber photometry^19^ (Fig. 5G).

Note that the width of the CCF, determined by the parameter *μ* (Eq. 16), is strictly a function of the parameters of the diffusion-less system (i.e., a function of the parameters of the local interaction between CINs and DA), and are independent of the diffusion coefficient, *D_u_*, that affects the speed of the wave. When *D_u_* is large, the wave is both faster and has a spatially broader interface. Conversely, when *D_u_* is small, the wave is slower and has a narrower interface. Thus, in this case of a single diffusion coefficient (*D_ν_* = *0*), these two effects (speed and width of interface) cancel out, causing the functional shape of the CCF to be the same for a given temporal delay *τ*, independently of *D_u_*.

### Turing instability can trigger traveling waves of CIN and DA activity

For *D_v_* > 0, numerical simulations show that the traveling waves are still the solution to the system, but their shape is not given by Eq. 10. Moreover, Eq. 11 is no longer valid and the reversal of the velocity of the traveling wave occurs elsewhere in the phase diagram (Fig. 5C). Interestingly, for the case of finite *D_v_*, there exists a parameter regime where the uniform solution *(u_1_,v_1_*) (Eq. 7a) loses its stability through a Turing instability (Appendix 1). Spatial patterns (of a particular spatial scale, determined by the parameters) form spontaneously and transiently and trigger a traveling wave of advancing *u* and receding *v*, i.e., (*u_3_,v_3_*) (Eq. 7b) wins out (Fig. 6A,B, Suppl. Fig. 4). As explained in Appendix 1, the parameter regime where this is possible (provided *D_u_*/*D_v_* is sufficiently small) is determined by the inequality *A_11_*>*0*, which is an entry in the stability matrix of the diffusion-less system in the vicinity of one of the fixed-points (Eq. 7A), and which translates into (Fig. 6C, yellow region):

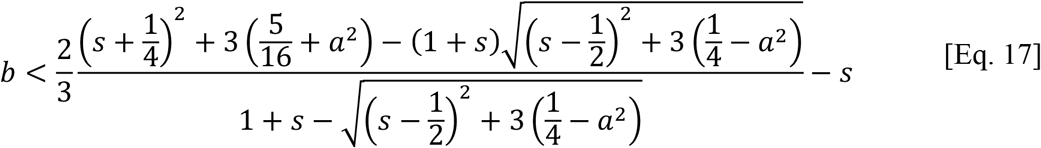

In Appendix 2, we analyze another parameter regime of the original model, which represents the case in which nAChRs on the GINs have a substantially higher affinity to ACh, than the nAChRs on the DA fibers. This regime can give rise to traveling waves of DA and CIN activity that advance together or recede together. Moreover, in a narrow parameter regime, receding waves that leave stable Turing patterns in their wake can also form.

**Figure 6.**
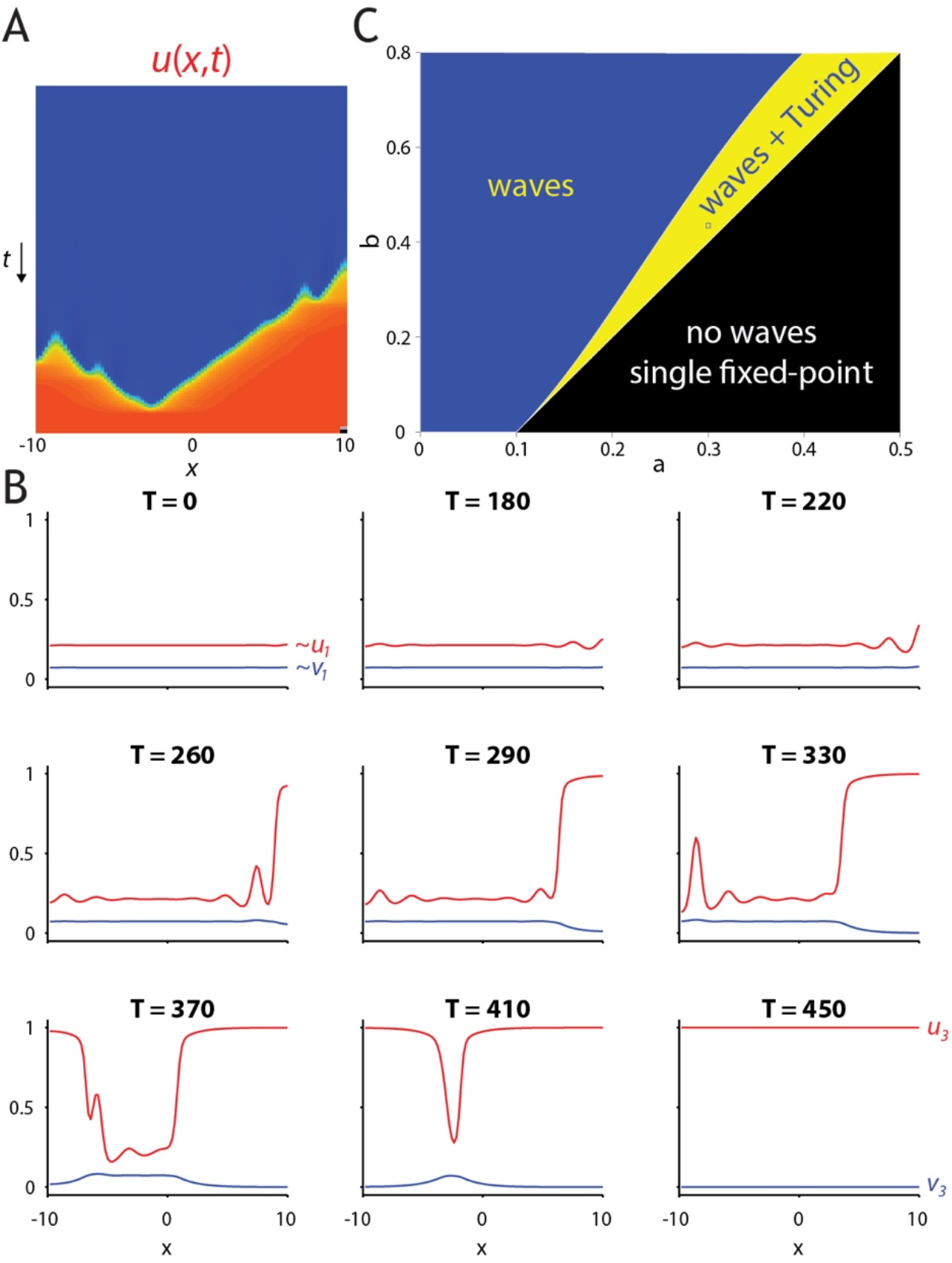
Turing instability in the tractable AIRD model. **A.** Space-time plot of *u*(*x,t*), and **B.**_ Series of snapshots describing the “spontaneous” destabilization by diffusion of the low *u* (CIN) / high *v* (DA) state, through a Turing instability. Transient localized “hills of activity” form, and they are overtaken by traveling waves that drive the system to the high *u* (CIN) / low *v* (DA) state. **C.** Indication of the region in the phase diagram where these transient Turing patterns can form (provided *D_u_/D_v_* is small enough). Parameters: *a=0.3, b=0.435, s=0.2, D_u_=0.06, D_v_=1*.

## Discussion

### Wavelike activity in the striatal cholinergic neuropil of freely moving mice

In this study, we have reported two novel experimental findings. First, implementing the method developed recently to visualize and analyze striatal DA waves^8^, we describe waves in the striatal cholinergic neuropil imaged using microendoscopy in freely moving mice. While we had shown preliminary hints of this in a previous publication, we now provide an analysis of these waves. Second, we demonstrated that the spatial extent of striatal DA release triggered with bipolar electrode stimulation depends on nAChRs that extend the range distance over which DA release occurs by several hundred micrometers, in agreement with a very recent study^20^.

Taken together these findings formed the basis for the formulation of the theoretical modeling part of this study. The fact that both DA and CIN activity form spatiotemporal wavelike activity, in conjunction with the fact the CINs can extend the spatial scale of DA release through nAChR activation, prompted us to hypothesize that it is the CINs that drive simultaneous waves of DA and CIN activity. We, therefore, investigated a common mechanism that could explain the existence of both spatial waves, based on the idea that the driver is the CINs. Moreover, the dynamic organization of these waves occurs locally in the striatum by direct interaction between CINs and DA fibers. A natural mechanism that fits these constraints is an AIRD system, in which the CINs are the activators and DA fibers are the inhibitors. Thus, we developed a dynamical model of coupling between CINs and DA fibers that relies on a physiologically-inspired model of their local interaction. We then supplemented this local interaction with a model of diffusion within a syncytium of cholinergic and dopaminergic neuropils. We demonstrated that this system can give rise to traveling wave solutions, and analyzed the parameters space of tractable systems that are essentially equivalent to the physiologically-inspired model, because they maintain the local geometry of nullclines and fixed-points.

### Nicotinic gain determines the direction of the traveling waves

The direction and velocity of the traveling wave solutions are determined by the parameters of the system. In our models, because of the bi-stability in the CIN dynamics, which gives rise to their putative self-excitation (see below), it is the CINs that drive the traveling waves, whereas the DA is the follower. Therefore, the strength of the CIN coupling to the DA fiber in the models is a critical parameter. In the tractable models, this is represented by the parameter *b* (which is roughly equivalent to the product of *γ* and *σ*, in the original physiologically-inspired model). When considering the phase diagrams of both models (Figs. 5D and A2.3C), it is evident that increasing *b* causes the direction of the traveling CIN wave to switch from advancing to receding. In the models where the expansion of CIN activity is coupled to the recession of DA activity (and vice versa), this makes sense, because increasing *b* – which signifies the increase of nicotinic gain to DA fibers – “strengthens the DA at the expense of the CINs”. Geometrically, increasing *b* (in both of the tractable models) increases the area between the nullclines *f_i_*(*u*) and *g_i_*(*u*) (Figs. 5A,C and A2.3A,B) from *u_1_* to *u_2_*, so that it eventually becomes larger than the area between these curves from *u_2_* to *u_3_* (Figs. 5A,C and A2.3A, B). This in turn, causes the wave to switch direction to where (*u_1_,v_1_*) wins out. This geometrical interpretations also helps to see why increasing *β* in the original model has the same effect (Figs. 4A,B). As *β* is increased, the area between the nullclines from *u_1_* to *u_2_* increases in the same fashion, pulling the traveling wave toward (*u_1_,v_1_*). *β* also represents nicotinic gain in the system. In contrast, it is the gain of the nAChRs on the GINs whose increase can cause the DA wave to advance in the original model, with *f_1_*(*u*) and *g_1_*(*u*), and in the tractable model with *f_2_*(*u*) and *g_2_*(*u*).

### Model assumptions: autocatalytic CINs and diffusion

The formation of wavefronts requires that the equation *f_i_*(*u*) (Eq. 6A) of the activator *u*(*x,t*) have an inverted-N shape, because the equation for the inhibitor *g_i_*(*u*) (Eq. 6B) can then intersect *f_i_*(*u*) at 3 points, thereby creating two stable fixed points (*u_1_,v_1_*) and (*u_3_,v_3_*). We posited that the CINs’ nullcline *f_i_*(*u*) inherits this shape from the known inverted-U shape of the dependence of nAChRs on ACh concentrations (which will be positively correlated with CIN activity). Because these nAChRs activate GINs^44, 54, 55^ which are inhibitory, the “negative sign” of inhibition contributes an upright-U shaped kink into the nullcline *f_i_*(*u*), creating its inverted-N shape (Figs. 4A,B; 5A,C & A2.3A,B). Physiologically, this means that for strong enough activation of CINs, the increase in ACh concentration (>1 mM, Fig. 3B) will lead to a de-activation of the nAChRs on the GINs, which would translate into a recurrent self-disinhibition of CINs by their own activation. This, in essence, is how we realized the self-excitatory (autocatalytic) nature of CINs in this model. If our proposed mechanism is to be taken seriously, one would need to demonstrate that α4β2 nAChRs on GINs indeed exhibit an inverted-U shape at physiological levels of striatal ACh, as was observed in *Xenopus* oocytes^47^. Similarly, in the models in which there is a high ACh/low DA fixed-point (*u_3_,ν_3_*), the slope of *f_i_*(*u*) at that fixed-point is negative. This means that the α4β2 nAChRs on the striatal DA fibers are also operating in the regime where high ACh deactivates them, which would also need to be demonstrated empirically. In the original model this required the value of *σ* to be close to 1, so that the inverted-U shape of *g_1_*(*u*) was visible in the same range of *u* values (i.e., CIN activation) where the inverted-N shape of *f_1_*(*u*) was visible (Fig. 4). Physiologically, this means that the affinity of the α4β2 nAChRs to ACh (which we are assuming is positively correlated with the CIN activity) is similar for both the DA fibers and the GINs. In contrast, when *σ* ≪ *1* (Appendix 2), which represents a case where the affinity of the nAChRs on the GINs is much higher than the affinity of the nAChRs on DA fibers, the kink in *f_1_*(*u*) occurs only in the rising part of *g*_1_(*u*), so that in this case CINs can only activate (but not deactivate) DA fibers.

The assumption that CINs can self-excite lies at the heart of the AIRD mechanism we propose for the generation of traveling waves. In the current state of experimental knowledge this assumption is quite tenuous. If anything, CINs are known to couple in acute striatal slice experiments through di-synaptic inhibition^39, 40^. However, it is possible that alternative mechanisms of self-excitation exist *in vivo* that could support our model and give rise to traveling waves. For example, DA fibers can co-release glutamate^56, 57, 58, 59, 60^. Activation of nAChRs on DA might lead to glutamate release which in turn would both excite CINs on a fast time scale through ionotropic glutamate receptors that would effectively provide self-excitation, and inhibit them more slowly by activation of metabotropic D2Rs. Alternatively, activation of presynaptic α7 nAChRs on cortical terminals might produce self-excitation of CINs. In such a scenario, activation of cortical input could introduce the bi-stability: when α7 nAChRs were activated, CINs would become autocatalytic because in this scenario their release of ACh could cause the cortical input to have a stronger excitatory influence over themselves causing self-excitation. This alternative scenario has the corollary that in the absence of cortical drive CINs cannot be autocatalytic. While all of these proposed mechanisms are speculative, they suggest predictions that can be tested experimentally, and the existence of DA and CIN waves warrants a search for some mechanistic explanation for their generation.

The diffusion term introduced into the model functions to enable the spread of activation of DA and CIN activity. We have based the diffusion component on the fact that both DA axons^6, 61, 62, 63, 64^ and CIN axons have release sites every few microns^38, 65, 66, 67, 68^ and are space filling, so they can be conceptualized as syncytia that interact everywhere, and that once an area is activated there is potential for that activation to spread. Physical diffusion of DA and ACh from release sites is too short-ranged to produce the spread of activation predicted by the model. On the other hand, the time scales of such electrical spread are on the order of milliseconds to 10s of milliseconds, which is probably faster than the spreading of wavelike activity observed by ourselves and others^8^. Of course, the slower observed dynamics may reflect the time constants of the decay of the fluorescence, and thus do not rule out electrotonic spread. With all these caveats in mind, we considered two cases: one in which *D_ν_* = *0*, which we interpret to represent a case where all the DA released is caused by CIN activation. The only activity that actually spreads is that of the CINs, and when a nAChRs is activated on a DA fiber it cannot trigger the spread of activity in the DA fibers but only trigger local release of DA. The second case we considered is that of *D_ν_*>*0* (we used, without loss of generality, *D_ν_*=*1*). Here the assumption is that activity can also spread in the DA fiber syncytium, and that generally the diffusion of this activity is more long-ranged (as discussed below).

### Wave propagation and temporal correlations between DA and CIN activity

In the absence of direct visualization of spatiotemporal neural activity patterns, calculating the CCF between two signals is a widely used method to characterize the temporal correlation between them. Using the tractable models (*i*=*2,3* in Eqs. 6), we derived the structure of the expected CCFs between the DA and CIN signals at a given point, *x_0_* [i.e. *v*(*x_0,t_*) and *u*(*x_0,t_*), respectively]. In the case when there is a single diffusion coefficient *D_u_* in the problem (i.e., *D_v_* = 0), the width of the CCF is independent of *D_u_* (Eqs. 16 and A2.5), meaning that even if there is a mismatch between the true underlying timescale of diffusion and the experimentally observed one, this mismatch should not strongly impact the measured temporal correlations. The tacit assumption in our calculation of the CCF is that our measurement is insensitive to the baseline value at a given region but only to the changes from that value (a reasonable assumption for many of the existing experimental techniques such as fiber photometry). Hence, the calculation of the CCF using the temporal derivatives of *U* and *V* (Eqs. 10 and A2.3). For the case where the two waves advanced or receded together (Appendix 2), we found that the correlation is symmetric and in phase. Symmetric, in-phase CCFs have typically not been found between DA and CIN signals in the basal ganglia, arguing against coupled DA and CIN waves that advance or recede together.

The tractable model in which one wave spreads while the other recedes can give rise to a richer variety of CCFs (Eq. 14). Importantly, the shape of the CCF depends on the direction of the two waves: a reversal in the wave direction reversed the temporal relationship between DA and CINs. Interestingly, the direction of the empirically observed DA waves in dorsal striatum is determined by the nature of the task: medial to lateral waves are associated with instrumental learning, while lateral to medial waves were associated with the reward delivery during classical conditioning^8^. Thus, a prediction of our model would be that each of these tasks should give rise to a different temporal correlation structure. Curiously, the relationship between CINs and midbrain DA neurons does indeed change depending on the nature of the cue, reward etc. ^29, 30^.

The direction of wave propagation required in the model to generate a CCF that resembles the empirically observed one (Figs. 5F,G)^19^ is such that the CINs wavefront is receding. The delayed peak DA activity relative to CIN activity, actually arises in the model from the drop in CIN activation preceding a drop in DA. This result is actually quite intriguing because the main behaviorally relevant signal of CINs is their famous pause response, in which their firing rate abruptly drops in response to reward or stimuli related to reward^29, 69, 70, 71, 72^.

### Turing mechanism

We found that the effects of a Turing bifurcation could be observed both in the regime where the fixed points were high DA/low CIN and vice versa (see above), and in the regime where the fixed points were high DA/high CIN and low DA/low CIN (Appendix 2). In the former case, the fixed point that lost stability was that of high DA/low CIN (e.g., Eq. 7a). In this scenario, while Turing patterns could not be stabilized, the instability could play a role in driving the spontaneous formation of traveling waves. In other words, if the striatum were in a state of low CIN activity, and something drove an elevated global level of DA (e.g., with input from the SNc) then that state would lose stability and a traveling wave would form that would drive CIN activity high and lower the DA levels (Fig. 6, Suppl. Fig. 4). In the latter case (Appendix 2), the fixed point that lost stability was that of high DA/high ACh. (e.g., Eq. A2.1b). In this scenario, a uniform elevation in both CIN and DA would lose stability, and a receding wave of both profiles would form but would leave transient or stable “hills of activity” with the hill of high DA being broader and engulfing the hill of CIN activity (Fig. A2.3E, Suppl. Figs. 6 & 8). We argued above that the CCFs in this regime gave rise to in-phase synchrony between CIN and DA activity (which remains true in the scenario where Turing patterns are formed, not shown, and) which does not agree with the experimental data. Nevertheless, this regime offers a method by which the striatum can undergo a dynamical parcellation or tiling into distinct functional modules of high and low neuromodulatory activity.

The existence of Turing bifurcations and patterns, requires that *D_u_*/*D_v_* ≪ *1* (Eq. A1.2). Is such a regime physiologically relevant? There are two considerations that argue that it may be. First, the axonal arbor of CINs has a radius, *I_CIN_*, of approximately 0.5 mm^73^ whereas the radius of the axonal arbor of an individual DA, *I_DA_*, neuron is approximately 1 mm^5, 57, 74^. Diffusion coefficients scale the like the square of their corresponding diffusion lengths. Therefore, we can expect *D_u_/D_v_* < (*l_CIN_/l_DA_*)^2^. Physiologically, this means that the axon potential propagation throughout the DA axon arbor will encompass a wider volume than that of the CIN axon arbor. Thus the DA axon arbor will inhibit CINs (via D2Rs) over a longer distance than the reciprocal excitation caused by CINs on DA axons via nAChRs^20^.

An alternative view to consider is that sequential recruitment of midbrain DA neurons might be the cause of traveling DA waves. Combined with a coherent mapping of the axonal arbors of these neurons in the striatum, sequential activity might give rise to the traveling waves^8, 75, 76, 77^. While this scenario is possible, there is no evidence for sequential firing activity of midbrain DA neurons. Also, if the source of the traveling wave is in the midbrain, then the question of mechanism of wave formation just moves one synapse back, and still needs to be explained. It is also questionable whether sequential firing activity at the soma level would reliably translate into waves in the striatum, due to the diverse shapes of DA arbors^5^. Thus, the hypothesis that DA waves are generated by sequential activity of midbrain dopamine neurons currently lacks support.

### Summary and central predictions

We have provided evidence for striatal waves of CIN activity (Fig. 1), and that nAChRs subserve long range striatal DA release (Fig. 2). Taken alongside the recent observation of striatal waves of DA^8^, our findings raise the likelihood that DA and CIN activity waves are formed by a local (i.e., striatal) mechanism of wave generation. We proposed a physiologically-plausible dynamic mechanism that invokes an AIRD system. The central predictions that arise from our study are that simultaneous imaging of DA and CIN activity should reveal that these waves are in fact strongly coupled both spatially and temporally, most likely, such that one advances while the other recedes. Moreover, our models do not invoke CIN synchrony to produce the waves and are more consistent with a scenario wherein an individual CIN can cause DA release, without depending on synchrony of CINs. While this assumption is at variance with the current experimental evidence^20, 21, 22^, we nevertheless predict that activation of an individual CIN should cause local release of DA from nearby axons. These predictions await experimental confirmation or refutation.

## Appendix 1: Turing instability

In our models, we consider situations where the presence of diffusion will cause one of the uniform fixed points of Eqs. 6 to become destabilized through a Turing bifurcation to form Turing patterns^53^. These are spatial activity patterns of a particular spatial scale that is determined by the parameters of the problem. For this analysis, we will need to consider the stability matrix, *A*, of Eqs. 6 which will have the form

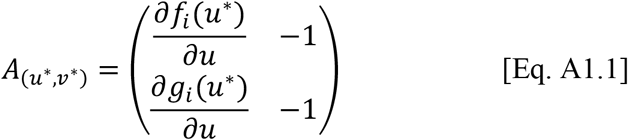

Where (*u*\*,*v*\*) is one of the stable fixed point of Eqs. 6 in the absence of diffusion. The conditions for the uniform solution (*u*\*,*v*\*) to lose stability through a Turing bifurcation are: a) *A_11_* > *0*; b) *A_12_A_21_* < *0*; c) trace of *A* is negative; and d)

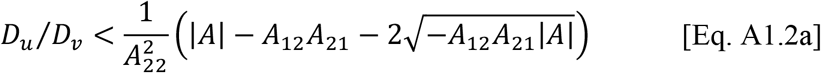

where |*A*| is the determinant of *A*^53^. Because *A_12_* = *A_22_* = −*1*, Eq. A1.2a simplifies to

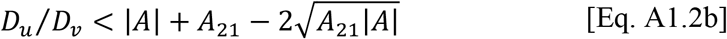

In the models that we analyze, calculations show that: the trace of *A* (i.e., 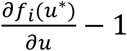) will be negative; 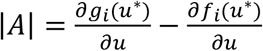 will be positive; and 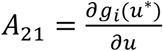 will be positive in the entire parameter regime of the candidate fixed-point (*u*,v\**) (the other stable fixed point cannot undergo a Turing bifurcation). Geometrically, this means that at the stable fixed point (*u*\*,*v*\*), that can lose stability through a Turing instability, the slope of *g_i_*(*u*) is always greater than the slope of *f_i_*(*u*), (that latter of which is always smaller than 1). Thus, by the inequality of arithmetic and geometric means, the right-hand-side of Eqs. A1.2 is always positive, so there will always be a small enough ratio *D_u_/D_v_* to attain a Turing bifurcation, provided *A_11_* > 0. Thus, in the models we will analyze, the parameter regime where Turing bifurcations are attainable is defined by the curve 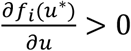 (which is indicated as yellow regions in the phase diagrams in Figs. 6C and A2.3C). Geometrically, what this boils down to, is that the slope of *f_i_*(*u*) at the Turing bifurcation point is positive, which is why *f_i_*(*u*) must have an inverted-N shape. Physiologically, this means that CINs must excite themselves, which we proposed could happen through recurrent-disinhibition, due to the putative U-shaped dependence on CIN activity of the nAChRs (located on the GINs).

## Appendix 2: Simultaneous advancing or receding traveling waves of CIN and DA activity

In the model we studied in the main text, while the wavefronts of ACh and DA overlapped and traveled together, one receded while the other advanced. This resulted from the geometry of the intersections between the nullclines which gave rise to two stable fixed-points of high *ν* (DA) and low *u* (ACh) or *ν* vice versa. In order for the two traveling waves to advance together or recede together the nullclines need to intersect such that they give rise to one fixed-point of low *u* and low *ν* and another of high *u* and high *ν*. This can be attained in the original model [with *f*_1_(*u*) and *g*_1_(*u*)] if *σ* ≪ 1 (Fig. A2.1). Physiologically, this represents the case where the nAChRs on the GINs have a substantially higher affinity to ACh, than the nAChRs on the DA fibers. As expected, for appropriate parameters the traveling waves of DA and ACh in this parameter regime advance or recede together (Fig. A2.1, Suppl. Fig. 5).

**Figure A2.1.**
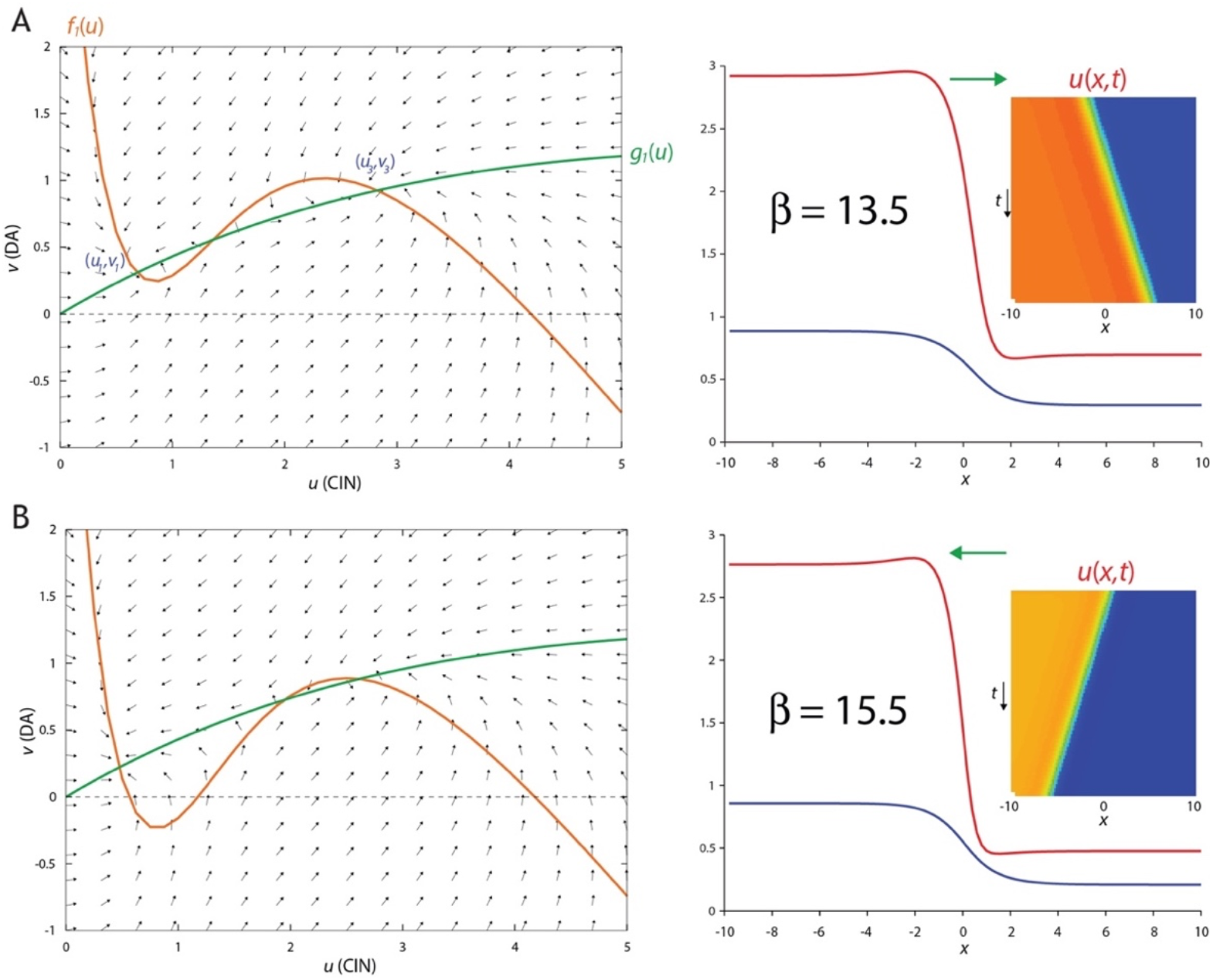
Simultaneously advancing or receding traveling waves in original model [with *f_1_*(*u*) and *g_1_*(*u*)]. **A**. Left: Phase plane. setting *σ*=0.1 horizontally stretches the green nullcline (which is equivalent to reducing the affinity of the nAChRs on the DA fibers to ACh) causing it to intersect the orange nullcline at three fixed-points, with the two stable fixed-points arranged such that *u_3_* > *u_1_* and *ν_3_* > *ν_1_*. When *β*=*13.5*, the area between the nullclines adjacent to (*u_3_,v_3_*) is larger than the area adjacent to (*u_1_,v_1_*). Right: *u* and *v* form traveling waves where both fronts advance (rightwards) together. Inset: space-time plot. **B.** Left: Same as A, except that setting *β=15.5* causes the area between the nullclines adjacent to (*u_1_,v_1_*) to be larger. Right: Same as A, except that now the fronts recede (leftwards) together. Other parameters:*A=4.3, κ=1.5, g=4.7, D_u_=0.2, D_v_=1*.

Interestingly, if we choose parameters where the stable high *u* and high *ν* fixed-point of the diffusion-less model occurs where the slope of *f*_1_(*u*) is positive (Fig. A2.2A), the model gives rise to receding traveling waves that leave a spatial pattern in their wake (Fig. A2.2B,C, Suppl. Fig. 6).

**Figure A2.2.**
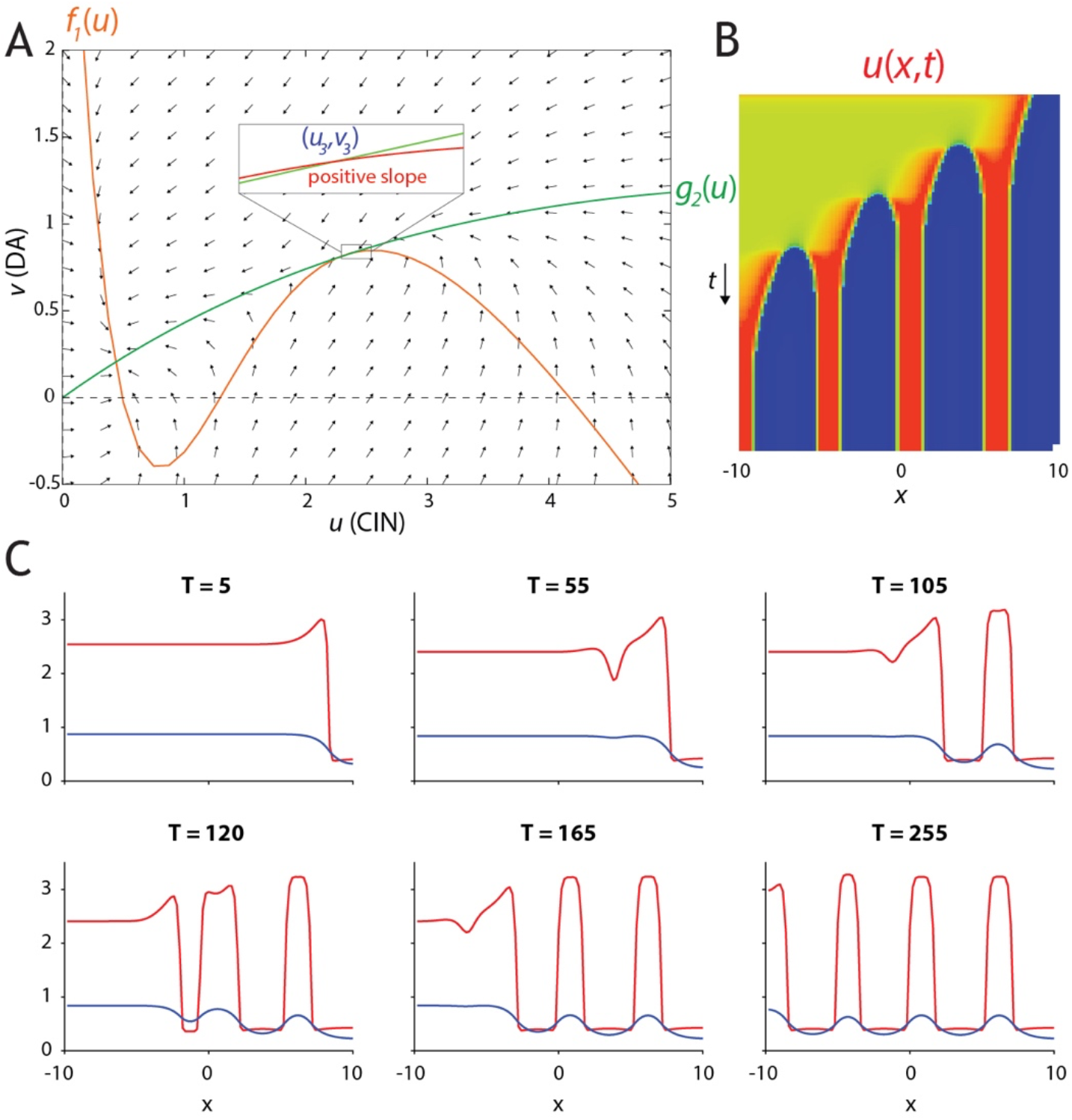
Stable Turing patterns form in the wake of receding traveling waves in original model [with *f_1_*(*u*) and *g_1_*(*u*)]. **A.** Phase plane. when *β=16.2* the area between the nullclines adjacent to (*u_3_,ν_3_*) is very small, the two nullclines cross with positive slopes (inset) that fulfill the condition for a stable Turing bifurcation (see Appendix 1). **B.** In this parameter regime, the space-time plots reveal a receding traveling wave that leaves stationary and localized peaks of *u* (and *ν*, not shown) standing **C.** Snapshots of the receding wavefronts reveal how they leave localized hills of activity in its wake, with the spatial spread of the *ν* (DA) being wider than that of *u* (CIN). Other parameters: *A=4.3, σ=0.1, κ=1.5, γ=5, D_u_=0.01, D_ν_=1*.

To gain a fuller, analytical understanding of this parameter regime and of the pattern formation, we once again replace *f*_1_(*u*) and *g*_1_(*u*) with tractable polynomial models: *f*_3_(*u*) = *u(1–u)(u–s)* and *g*_3_(*u*) = *bu*, with *0*<*s*<*1* and *b*>*0*. This model (Fig. A2.3) is known as the Fitzhugh-Nagumo model^78^.

In the absence of diffusion, this system has two stable fixed points:

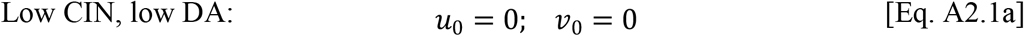

and

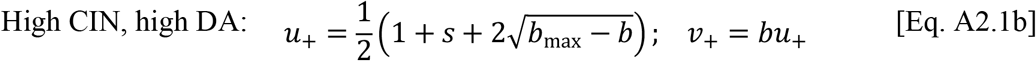

with *b_max_* ≝ (1 – *s*)^2^/4, so that the condition for the existence of the second stable fixed point is that *b* ≤ *b*_max_.

**Figure A2.3.**
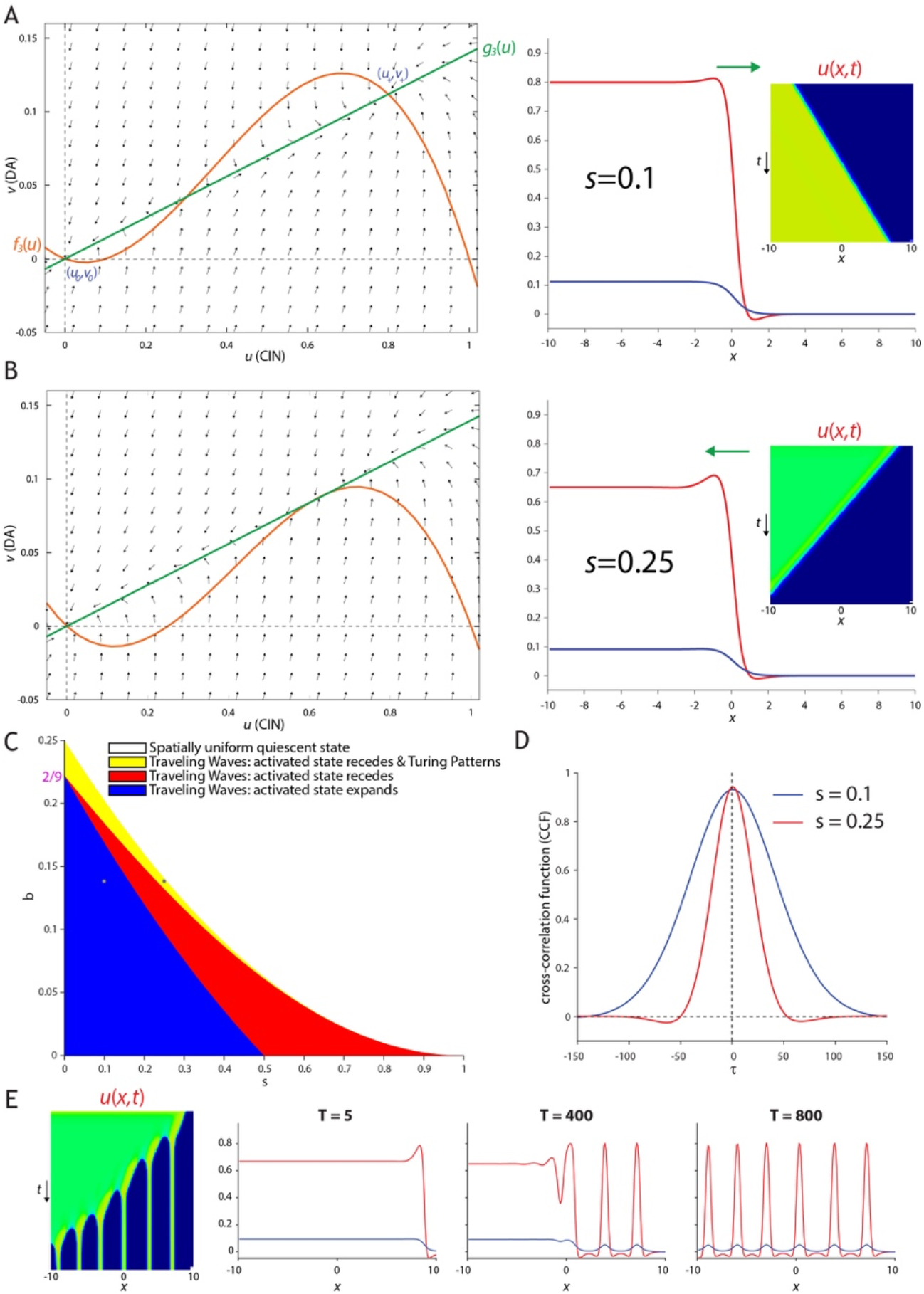
Fitzhugh-Nagumo (FN) model for simultaneous advancing and receding traveling waves and Turing patterns [with *f_3_*(*u*) and *g_3_*(*u*)]. **A.** Phase plane. Arrangement (Left) of fixed-points (*e.g., u*_+_ > *u_0_* and *ν_+_* > *ν_0_*) and nullclines [*e.g*., area between nullclines adjacent to (*u+,v+*) is larger] that give rise to simultaneously expanding “activated” states (Right). **B.** Same as A, except that area between nullclines adjacent to (*u_0_,ν_0_*) is larger, giving rise simultaneously receding activated states Insets: space-time plots. **C.** Phase diagram of FN model, demonstrates that it supports both advancing and receding waves, and in a subset of the receding wave regime it is possible to observe Turing patterns (provided the *D_u_/D_ν_* is sufficiently small). **D.** Cross-correlation function [of *u*(*x_0,t_*) and *v*(*x_0,t_*)] during a traveling wave is always symmetrical. Parameters correspond to black and yellow asterisks in C. **E.** Space-time plot and snapshots of a receding wave leaving Turing patterns in its wave. Parameters correspond to black asterisk in C.

Simulating the system with diffusion (i.e., *D_u_ = 0.1* and *D_v_ = 1*) demonstrated that the two wavefronts advanced or receded together (Figs. A2.3A,B, Suppl. Fig. 7). Repeating the traveling wave analysis (Eqs. 8) for this model, with *D_v_ = 0*, we find traveling wave solution whose velocity is given by

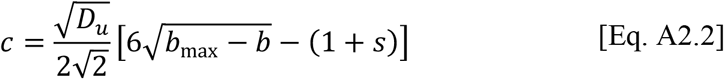

For the case where *c* = 0, we can derive a precise waveform solution for the standing wave

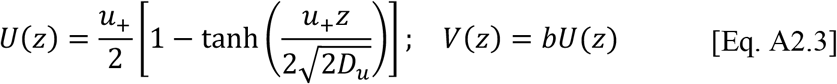

The condition *c = 0*, can be also derived from the requirement 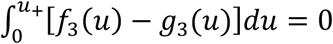 which yields

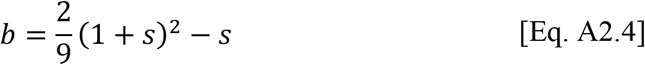

This curve (Eq. A2.4) divides the phase diagram into two regions: one where the waveforms for *u* and *ν* recede (Fig. A2.3C, red region), and one were they advance (Fig. A2.3C, blue region). Calculation of CCF for Eqs. A2.3 yields a positive symmetric *C*(*τ*) = *C_S_*(*λτ*) (see Eqs. 15a) with

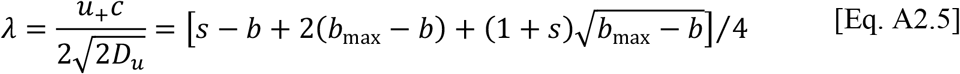

The symmetric CCF is maintained also when *D_ν_* > 0 (Fig. A2.3D). Thus, this model only gives rise to in-phase synchrony between CIN and DA. The width of the CCF depends on the velocity of the waves (Suppl. Fig. 7): when the wave is faster (e.g., for *s* = *0.25*), the CCF is narrower (Fig. A2.3D).

In the model discussed in the main text, while the Turing instability could give rise to transient spatial patterns it was impossible for these patterns to stabilize, because the two variables opposed each other at every given point: where one tended to a low value the other tended to a high value and vice versa, so this drove them apart and dissipated the initial destabilizing transient spatial pattern. In contrast, in the current model, where the two variables tend to high and low values together, it is possible that Turing patterns will stabilize with the “long-range” inhibition by DA separating space into localized regions that are elevated by the “short-range” excitation by CINs.

Conducting the stability analysis near the activated state (Eq. A2.1b), the condition for a Turing instability (provided is Eq. A1.2 is fulfilled) is (Fig. A2.3C, yellow region)

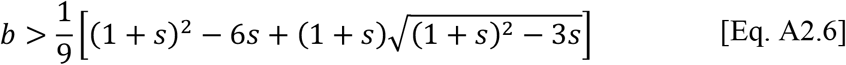

As expected, in this parameter regime we can observe receding waves that leave either transient or stable Turing patterns of localized activity (Fig. A2.3E, Suppl. Fig. 8). This analysis also provides an explanation for the patterns observed in the original model (Fig. A2.2B,C, Suppl. Fig. 6). In particular, we found that the patterns only occur in the original model [*f_1_*(*u*), *g_1_*(*u*)] when the nullclines intersect such that unstable fixed point is close to the fixed point with the high values of *u_3_* and *ν_3_*. This corresponds to the yellow band in Fig. A2.3C in the tractable model *f_3_*(*u*), *g_3_*(*u*)] being adjacent to the region where that two fixed points are close to merging – which is close to the region where only a uniform trivial solution exists – namely when *b* is close to *b*_max_. In these narrow regions there can occur a stable fixed point (*u*_+_, *ν*_+_) where the slopes of *f_1_*(*u*) (Fig. A2.2A) or *f_3_*(*u*) are still positive.

## Acknowledgments

This work was funded by a Research Grant from the Human Frontier Science Program (HFSP) no. RGP0062/2019 to J.R.W. and J.A.G, and by an ERC Consolidator Grant (no. 646886) to J.A.G.

## Author Contributions

Conceptualization, J.R.W. & J.A.G.; Methodology, L.M., N.G., Y.A., L.T. & J.A.G.; Software, L.M., N.G., Y.A., L.T. & J.A.G.; Validation, L.M., N.G., Y.A., L.T. & J.A.G.; Analysis, L.M., N.G., Y.A., L.T. & J.A.G.; Resources, J.A.G.; Data Curation, L.M., N.G., Y.A., L.T. & J.A.G.; Writing, L.M., N.G., Y.A, J.R.W. & J.A.G.; Visualization, L.M., N.G. & J.A.G.; Supervision, J.A.G.; Project Administration and Funding Acquisition, J.R.W. & J.A.G.

## Methods

### Animals

Experimental procedures adhered to and received prior written approval from the Hebrew University’s Institutional Animal Care and Use Committee. Two-to-nine-months-old male choline acetyltransferase-cre dependent (ChAT-IRES-Cre) transgenic mice (stock number 006410; Jackson Laboratories, Bar Harbor, ME, USA) were used for the microendoscopic imaging experiments. Choline acetyltransferase-cre dependent (ChAT-IRES-Cre) transgenic mice (stock number 031661; Jackson Laboratories, Bar Harbor, ME, USA) were cross-bred with Cre-dependent, Tet-controllable, GCaMP6f (Ai148, TIT2L-GC6f-ICL-tTA2) mice (stock number 030328; Jackson Laboratories, Bar Harbor, ME, USA) to create a double transgenic mouse line expressing GCaMP6f in cholinergic cells. Two-to-seven-months-old double transgenic mice or wildtype C57BL/6J mice, that underwent stereotaxic viral inoculation, of both sexes were used for electrical stimulation experiments. All mice were on a C57BL/6J background and housed under a 12-h light/dark cycle with food and water *ad libitum*.

### Stereotaxic Injection of Adeno-Associated Virus

Mice were deeply anesthetized with isoflurane in a non-rebreathing system (2.5% induction, 1–1.5% maintenance) and placed in a stereotaxic frame (model 940, Kopf Instruments, Tujunga, CA, USA). Temperature was maintained at 35°C with a heating pad, artificial tears were applied to prevent corneal drying, and animals were hydrated with a bolus of injectable saline (5 ml/kg) mixed with an analgesic (5 mg/kg carprofen or meloxicam). To calibrate specific injection coordinates, the distance between bregma and lambda bones was measured and stereotaxic placement of the mice was readjusted to achieve maximum accuracy. A small hole was bored into the skull with a micro drill bit and a glass pipette was slowly inserted at the relevant coordinates in aseptic conditions.

A total amount of 400 nl of an adeno-associated virus serotype 9 harboring GCaMP6s construct (AAV9-syn-flex-GCaMP6s; > 2.5×10^13^ viral genome/ml; University of Pennsylvania Vector Core, catalog number AV-9-PV2824) was manually injected into the dorsal striatum under aseptic conditions. The coordinates of the injection were as follows: anteroposterior, +0.5 mm; mediolateral, +2.3 mm; and dorsoventral, −2.8 mm, relative to bregma using a flat skull position^79^.

A total amount of 250 nl of an adeno-associated virus serotype 5 harboring GRAB-DA2m construct (AAV5-hSyn-DA2m; > 4.85×10^13^ VG/ml viral genome/ml; WZ biosciences inc Lot No. 20210119) was injected with the *Nanoject III* system (Drummond) into the substantia nigra under aseptic conditions. The coordinates of the injection were as follows: anteroposterior, −3.1 mm; mediolateral, +1.2 mm; and dorsoventral, −4.2 mm, relative to bregma using a flat skull position. To minimize backflow, solution was slowly injected and the pipette was left in place for 5-7 min before slowly retracting it.

### Gradient Refractive Index (GRIN) Lens Implantation

One week after the stereotaxic injection, mice were deeply anesthetized with isoflurane in a non-rebreathing system and placed in the stereotaxic frame, as described above (in some cases, a bolus of ketamine (32 mg/kg)-xylazine (3.2 mg/kg) was injected initially to stabilize the preparation for and induction of anesthesia). A hole slightly wider than the 1mm diameter (4 mm long) GRIN lens was drilled into the skull in aseptic conditions. We aspirated cortex with a 27-30 Gauge needle to a depth of approximately 2 mm (just past the corpus callosum) and then fit the lens in snugly and (dental-) cemented it into place together with a head bar for restraining the mouse when necessary. One week later we attached a baseplate to guarantee that the endoscope will be properly aligned with the implanted GRIN lens. To ensure light impermeability, the dental cement was mixed with coal powder and painted with black nail polish.

### Microendoscopic Imaging

Microendoscopes (nVista, Inscopix, Palo Alto, CA, USA) sampled an area of ~600 × 900 μm (pixel dimension: 1.2 μm) at 10 frames/s. Movies were motion corrected with the Inscopix Data Processing Software suite (IDPS, Inscopix, Palo Alto, CA, USA). The constrained non-negative matrix factorization (CNMF-E) algorithm (Zhou et al 2018, Pnevmatikakis et al 2015) was applied to motion corrected movies to detect putative somata and to estimate the background signal. Fluorescent movies of the background signal alone were obtained by the CNMF-E algorithm and fluorescence changes over time (Δ*F/F_0_*) were extracted such that 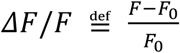, where *F* is the raw fluorescent values recorded; *F_0_* denotes the minimal averaged fluorescence in 1.5 sec-overlapping 3 sec periods throughout the measurement.

### Behavior

Two weeks after the lens implantation the nVista microendoscope was attached to the baseplate and mice were placed for imaging in a behavior chamber (21 cm × 13 cm × 8 cm) lit by diffuse infrared light. Mice were free to move around the behavior chamber during the imaging session.

### Acute slice preparation

Two to three weeks after the viral injections mice were deeply anesthetized with ketamine (200 mg/kg)–xylazine (23.32 mg/kg) and perfused transcardially with ice-cold modified artificial cerebrospinal fluid (ACSF) bubbled with 95% O_2_–5% CO_2_, and containing (in mM): 2.5 KCl, 26 NaHCO3, 1.25 Na_2_HPO_4_, 0.5 CaCl_2_, 10 MgSO_4_, 0.4 ascorbic acid, 10 glucose, and 210 sucrose. The brain was removed and sagittal slices sectioned at a thickness of 275 μm were obtained in ice-cold modified ACSF. Slices were then submerged in ACSF, bubbled with 95% O_2_–5% CO_2_, and containing (in mM): 2.5 KCl, 126 NaCl, 26 NaHCO_3_, 1.25 Na_2_HPO_4_, 2 CaCl_2_, 2 MgSO_4_, and 10 glucose, and stored at room temperature for at least 1 h prior to recording.

### Electrical stimulation protocol

Electrical stimulation was carried out by a parallel bipolar Platinum-Iridium electrode with diameter of 125 μm and spacing of 500 μm (FHC, PBSA0575). The magnitude of the stimulus was controlled through a stimulus isolator (ISO-Flex, MicroProbes) while the frequency and duration were controlled by computer software (MESc, Femtonix). A total of 10 pulses (10Hz, 2ms duration, 3mA) were delivered.

### Slice visualization, data collection and fitting – 2PLSM

The slices were transferred to the recording chamber of Femto2D-resonant scanner multiphoton system (Femtonics Ltd., Budapest, Hungary) and perfused with oxygenated ACSF at room temperature. A 16X, 0.8 NA water immersion objective was used to examine the slice using oblique illumination.

2PLSM Ca^2+^/monoamine imaging: The 2PLSM excitation source was a Chameleon Vision 2 tunable pulsed laser system (680–1,080 nm; Coherent Laser Group, Santa Clara, CA). Optical imaging of GCaMP6f/GRAB-DA2m signals was performed by using a 920-nm excitation beam. The fluorescence emission was detected and collected with gated GaAsP photomultiplier tubes (PMTs) for high sensitivity detection of fluorescence photons as part of the Femto2D-resonant scanner. The laser light transmitted through the sample was collected by the condenser lens and sent to another PMT to provide a bright-field transmission image in registration with the fluorescent images. Areas of ~340 μm × 340 μm were selected and imaged with 100 μm intervals from the electric stimulation center, ~31 Hz scans were performed, using ~0.6 μm pixels. ROIs were marked manually offline based on the online fluorescent traces that were extracted.

Optical and electrophysiological data were obtained using the software package MESc (Femtonics), which also integrates the control of all hardware units in the microscope. The software automates and synchronizes the imaging signals and electrophysiological protocols. Data was extracted from the MESc package (Femtonics) to personal computers using custom-made code in MATLAB (MathWorks, Natick, MA, USA) code. Fluorescent changes over time (Δ*F/F_0_*) datapoints were extracted such that 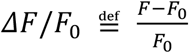, where *F* is the maximal fluorescent value recorded while evoking electrical stimulation; *F_0_* denotes the averaged fluorescence in the first second (pre-stimulation).

### Statistical Analysis

#### Endoscopic imaging

Regions in the fluorescent movies that were outside the GRIN lens were excluded from the analysis. The spatial average of the fluorescent field-of-view, was calculated, and the peaks and troughs of a filtered version, were extracted with custom-code. Amplitudes of each upswing (trough to next peak) that was larger than the average amplitude of upswings were used for the analysis.

Optic Flow analysis of the microendoscopic images was conducted as published^8^. The output of the analysis generates a vector field that changes with time (realized by assigning to each pixel at each time point a complex number whose modulus is the size of the vector and whose argument is its direction). In Fig. 1 & Suppl. Movie 1, we present a temporally filtered (7 frames) and spatially decimated (4 pixels) version of the vector field. To align the temporally filtered vector field with the original sequence of images, it was necessary to shift the vector field by approximately 4 frames in order to compensate for the shift generated by the temporal filtering.

For each upswing of the spatially averaged fluorescent signal, a spatial average of the flow field was calculated, after removing pixels whose vectors’ amplitudes did not exceed the 85^th^ percentile calculated using bootstrapping, as explained presently. If for a certain movie we identified *N* upswings in the average fluorescent signal, we calculated the mean length *L* (in frames) of these upswings. We then chose 1000 random frames. For each random frame we averaged the vector field over *L* frames following it, and created an averaged frame whose every pixel is represented by the modulus of the resulting vector. We then ordered for each pixel individually the 1000 moduli that arose from each of the repetitions. Then we set a threshold according to the 850^th^ largest modulus. We considered the vectors calculated for the upswing as significant if its modulus exceeded the threshold associated with its pixel.

#### 2PLSM resonant scanning

In Fig. 2C, Δ*F/F_0_* data points were fitted with a Lorentzian function 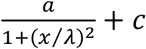 to extract the length scale *λ* of the fluorescence signals’ decay with distance. ANCOVA was used to test the hypothesis that the drop off of fluorescence with distance was significantly different in mecamylamine relative to control.

## KEY RESOURCES TABLE

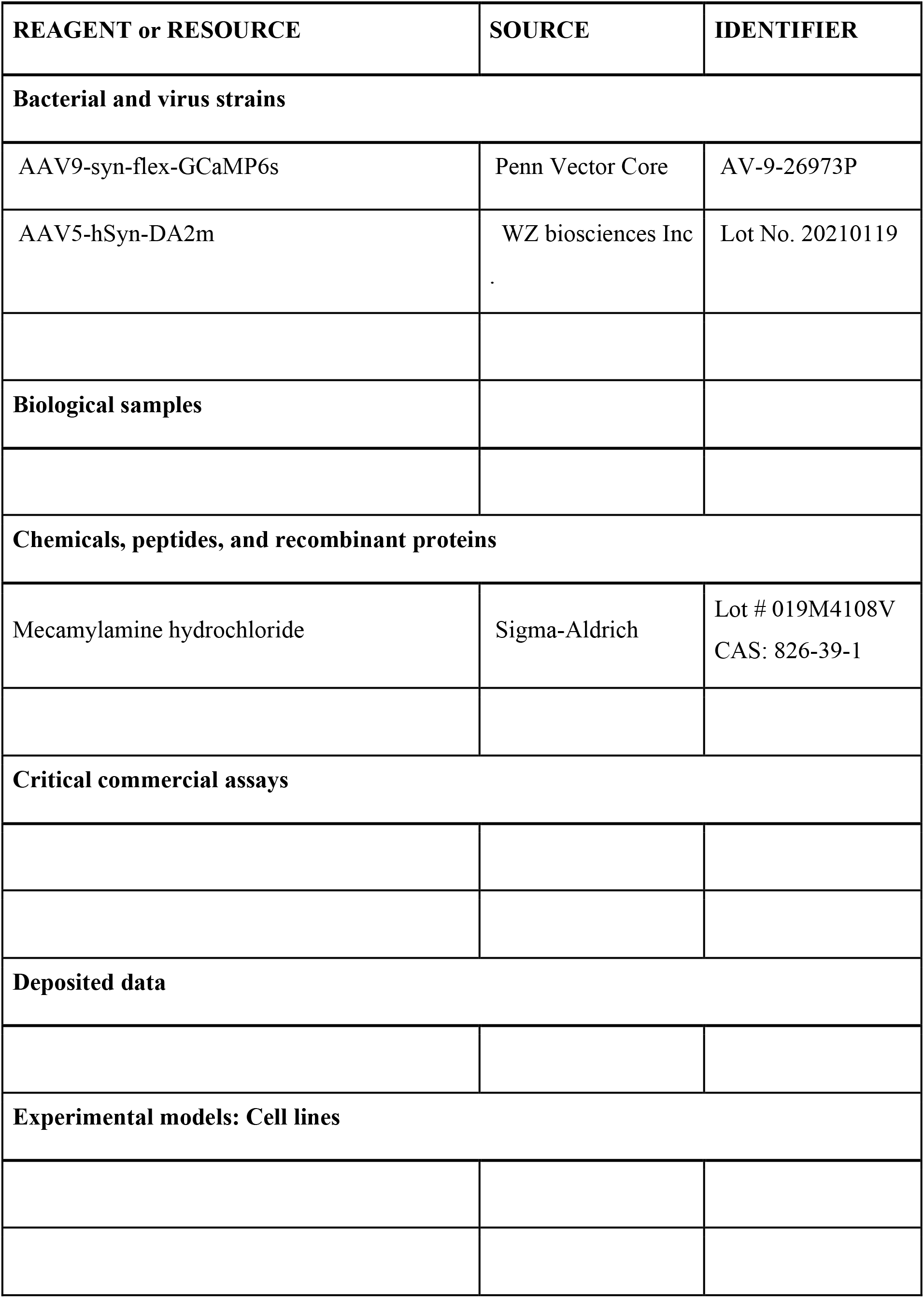
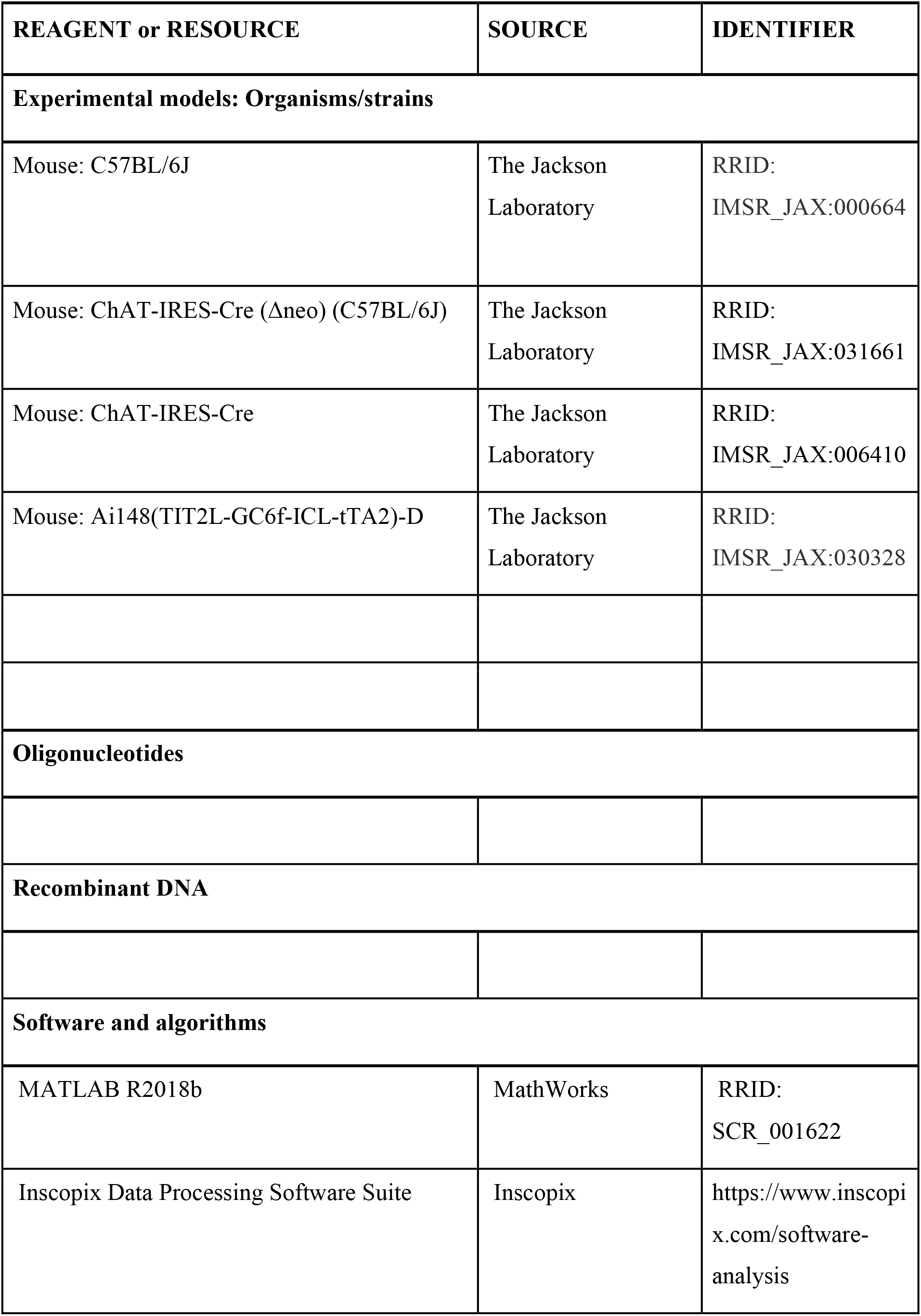
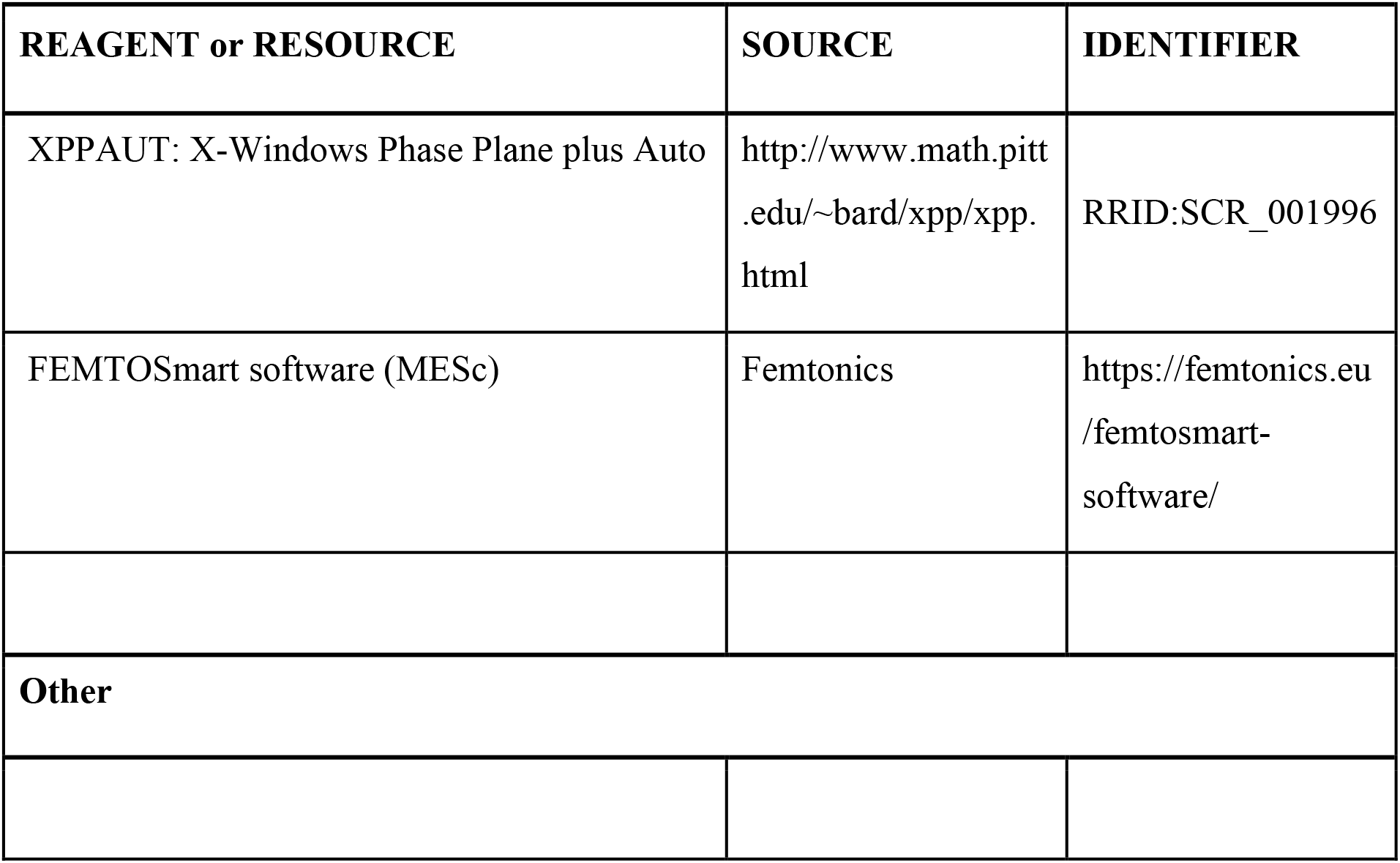

## Data and code availability

Any additional information required to re-analyze the data reported in this manuscript is available from the lead contact upon request. All the information necessary to run the XPPAUT simulations is present in the paper. Nevertheless, the lead contact will share code and simulated data files upon request.

