## Supplementary material for "Mechanism of dopamine traveling waves in the striatum: theory and experiment": Suppl. Fig. 2

| 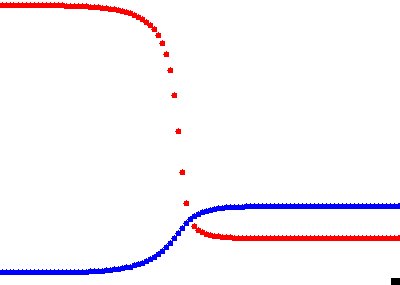 | 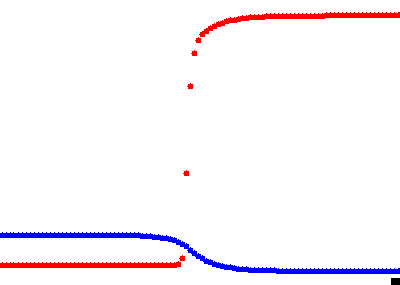 |
| --- | --- |
| *β=10, D_u_=0.02* (T=50) | *β=18, D_u_=0.02* (T=50) |
| 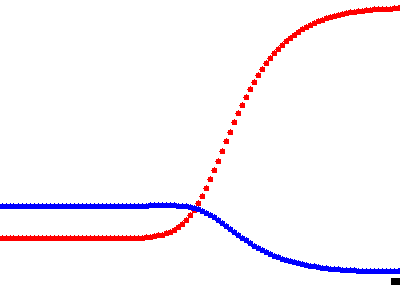 | 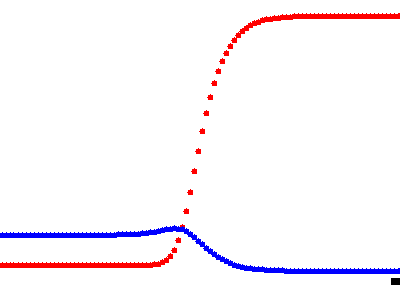 |
| *β=10, D_u_=1* (T=10) | *β=18, D_u_=1* (T=10) |
| **Supplementary Fig. 2A**: Original model with *f_1_*(*u*) and *g_1_*(*u*). red: *u*(*x,t*) (CIN); blue: *v*(*x,t*) (DA). Other parameters: *A=4.2, σ=0.75, κ=1.5, γ=4.7, D_v_=1*. Length of simulation in time units is indicated in parentheses. | |

| 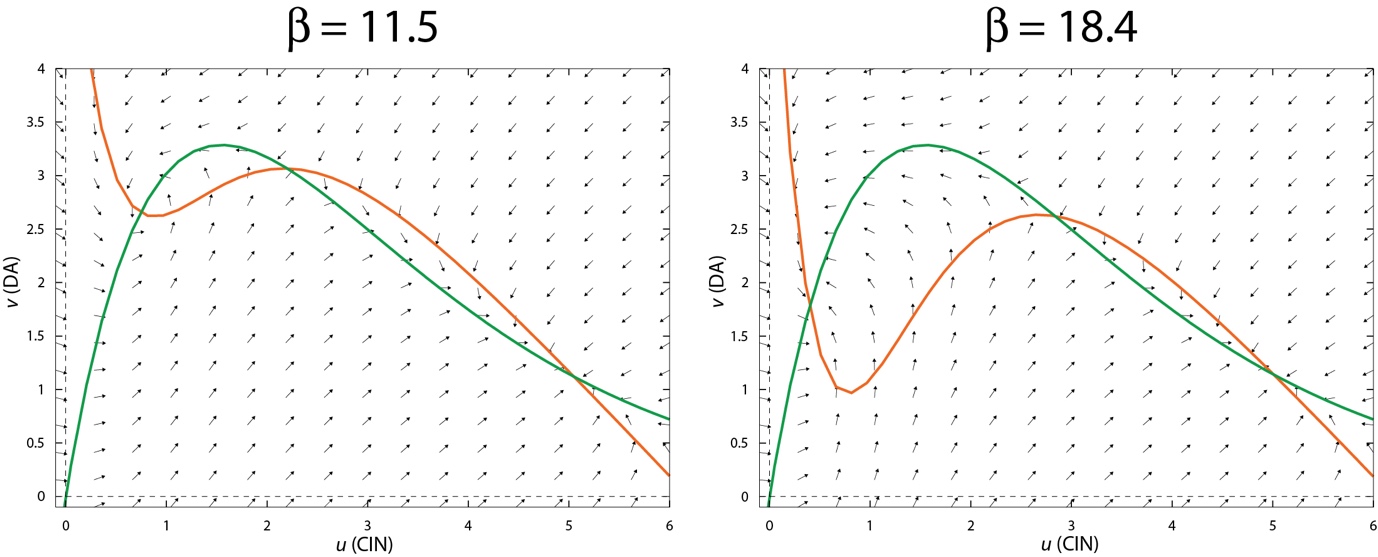 | |
| --- | --- |
| 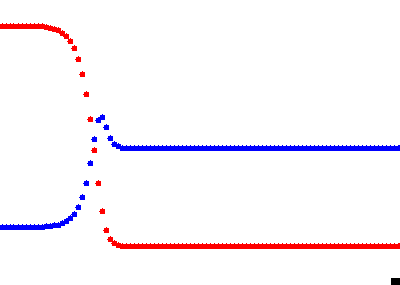 | 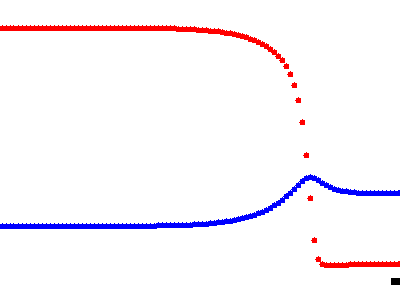 |
| *β=11.5*, *D_v_=0* (T=300) | *β=18.4, D_v_=0* (T=100) |
| **Supplementary Fig. 2B**: Original model with *f_1_*(*u*) and *g_1_*(*u*) with *D_v_=0*. red: *u*(*x,t*) (CIN); blue: *v*(*x,t*) (DA). Other parameters: A*=6.2, σ=0.43, κ=1.5, γ=13.4, D_u_=0.1*. Length of simulation in time units is indicated in parentheses. | |
