## Supplementary material for "Mechanism of dopamine traveling waves in the striatum: theory and experiment": Suppl. Fig. 3

| 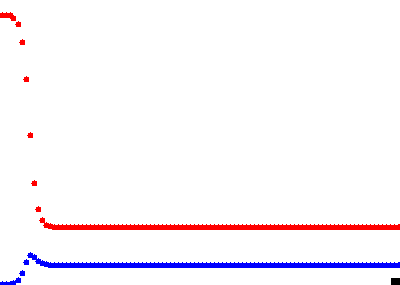 | 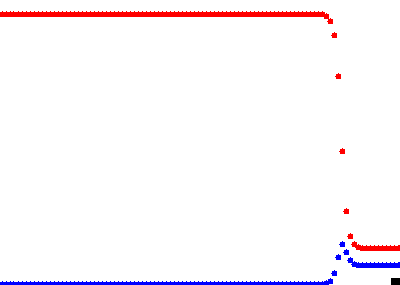 |
| --- | --- |
| *b=0.435* | *b=0.7* |
| **Supplementary Fig. 3**: Tractable model with *f_2_*(*u*) and *g_2_*(*u*) and *D_v_=0*. red: *u*(*x,t*) (CIN); blue: *v*(*x,t*) (DA). Other parameters: *a=0.3*, *s=0.2, D_u_=0.01.* | |
