## Supplementary material for "Mechanism of dopamine traveling waves in the striatum: theory and experiment": Suppl. Fi.g 4

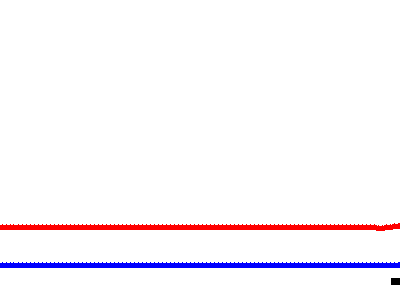

**Supplementary Fig. 4**: Turing instability in tractable model with *f_2_*(*u*) and *g_2_*(*u*). red: *u*(*x,t*) (CIN); blue: *v*(*x,t*) (DA). Other parameters: *a=0.3*, *b=0.435*, *s=0.2, D_u_=0.06*, *D_v_=1 .*
