## Supplementary material for "Mechanism of dopamine traveling waves in the striatum: theory and experiment": Suppl. Fig. 5

| 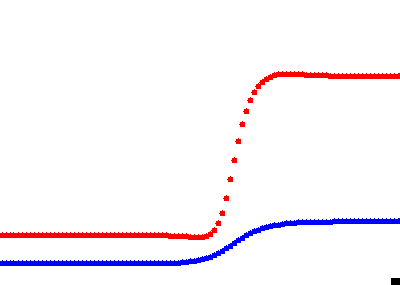 | 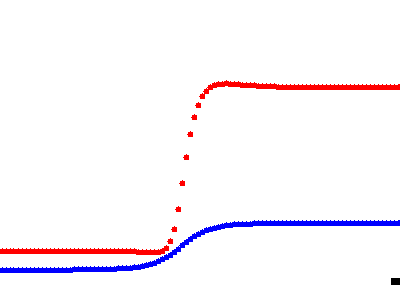 |
| --- | --- |
| *β=13.5* | *β=15.5* |
| **Supplementary Fig 5.** Full model with *f_1_*(*u*) and *g_1_*(*u*) with *σ=0.1*. red: *u*(*x,t*) (CIN); blue: *v*(*x,t*) (DA). Other parameters: *A=4.3, κ=1.5, γ=4.7, D_u_=0.2, D_v_=1*. | |
