## Supplementary material for "Mechanism of dopamine traveling waves in the striatum: theory and experiment": Suppl. Fig. 6

| 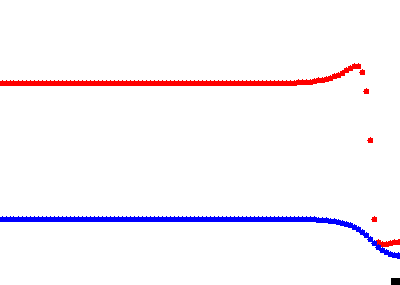 | 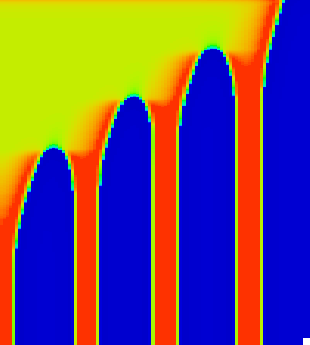 |
| --- | --- |
| *D_u_=0.01* | |
| 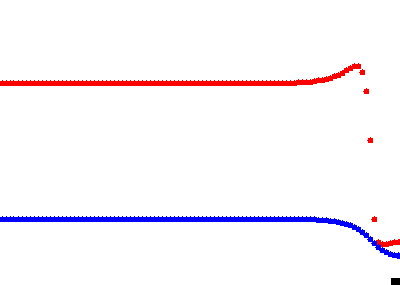 | 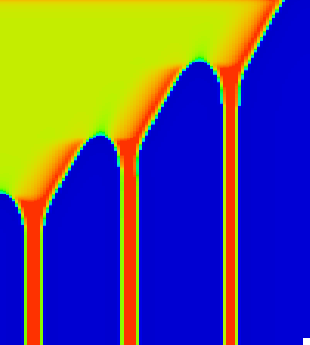 |
| *D_u_=0.015* | |
| 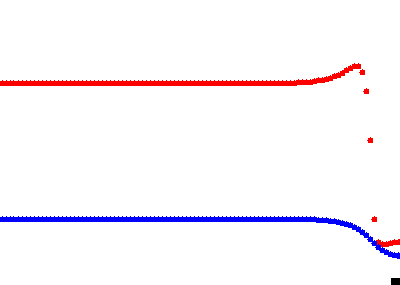 | 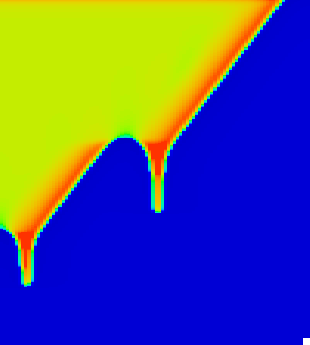 |
| *D_u_=0.02* | |
| **Supplementary Fig. 6**: Stable and unstable Turing patterns in full model with *f_1_*(*u*) and *g_1_*(*u*). red: *u*(*x,t*) (CIN); blue: *v*(*x,t*) (DA). Other parameters: *A=4.3, β=16.2, σ=0.1, κ=1.5, γ=5, D_v_=1*. | |
