## Supplementary material for "Mechanism of dopamine traveling waves in the striatum: theory and experiment": Suppl. Fig. 7

| 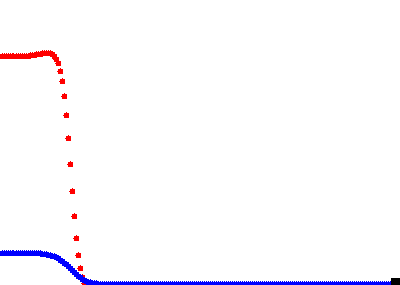 | 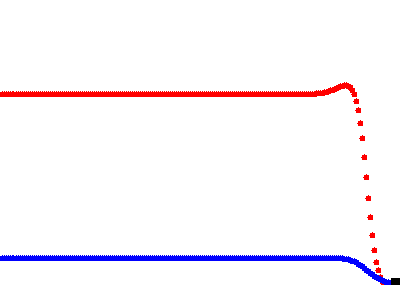 |
| --- | --- |
| *s=0.1* | *s=0.25* |
| **Supplementary Fig 7.** Tractable model with *f_3_*(*u*) and *g_3_*(*u*). red: *u*(*x,t*) (CIN); blue: *v*(*x,t*) (DA). Other parameters: *b=0.14, D_u_=0.1, D_v_=1*. | |
