## Supplementary material for "Mechanism of dopamine traveling waves in the striatum: theory and experiment": Suppl. Fig. 8

| 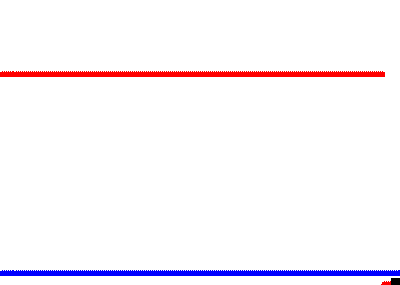 | 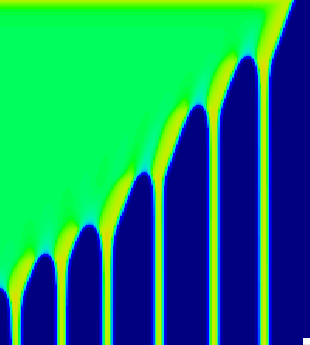 |
| --- | --- |
| *D_u_=0.02* | |
| 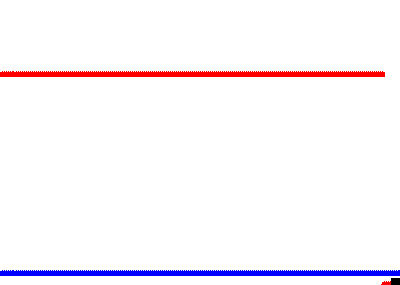 | 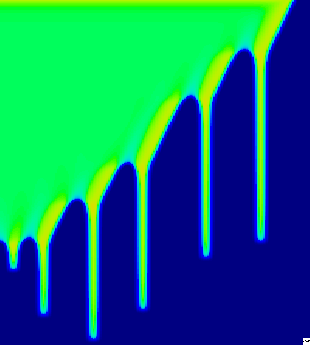 |
| *D_u_=0.022* | |
| 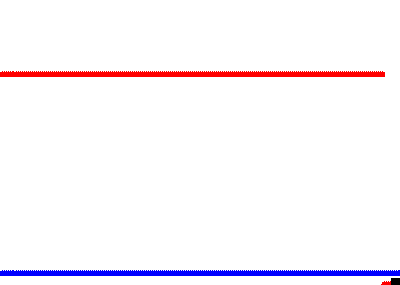 | 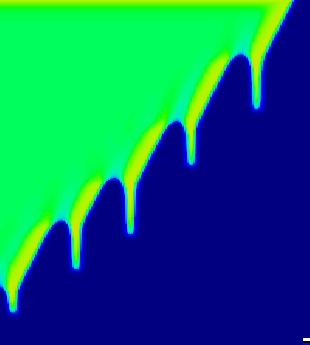 |
| *D_u_=0.024* | |
| **Supplementary Fig. 8**: Stable and unstable Turing patterns in tractable model with *f_3_*(*u*) and *g_3_*(*u*). red: *u*(*x,t*) (CIN); blue: *v*(*x,t*) (DA). Other parameters: *s=0.25 b=0.14, D_v_=1*. | |
